# Rice *Jumonji706* confers the photoperiod sensitivity in rice by distinct regulation of short-day and long-day flowering time regulatory pathways

**DOI:** 10.64898/2026.03.08.710421

**Authors:** Asanga D. Nagalla, Ryouhei Morita, Hiroyuki Ichida, Yoriko Hayashi, Yuki Shirakawa, Katsunori Ichinose, Tadashi Sato, Kinya Toriyama, Tomoko Abe

**Affiliations:** RIKEN Nishina Center for Accelerator-Based Science, 2-1 Hirosawa, Wako, Saitama 351-0198, Japan; Graduate School of Agricultural Science, Tohoku University, 468-1 Aramaki Aza Aoba, Aoba-ku, Sendai, Miyagi, 980-8572, Japan

## Abstract

Photoperiod sensitivity (PS) is a key biological response in plants as they adapt to specific environments. Rice (*Oryza sativa* L.) exhibits a clear PS, as it implements critical phase transition decisions based on PS signals. In this study, we identified a novel PS gene, *JMJ706,* that is expected to deliver photoperiod-related signals to the flowering-time regulatory network in a day-length-dependent manner. The *JMJ706* mutants exhibit early flowering under LD and later flowering under SD compared to WT plants. The gene encodes an H3K9me2 demethylase, and under long-day (LD) conditions, its demethylase activity facilitates the expression of *Grain number, Plant height, and Heading-date7* (*Ghd7*). Since *Ghd7* is a floral repressor in LD, it promotes the vegetative phase by delaying flowering. Under short-day conditions (SD), H3K9me2 demethylase activity facilitates *Early heading-date 1* (*Ehd1*) expression, and it acts as a floral accelerator by inducing *Heading date 3* (*Hd3a*) and *RICE FLOWERING LOCUS T 1* (*RFT1*). Furthermore, we propose that the daylength-dependent promotion of target genes (*Ghd7* and *Ehd1*) occurs through demethylation of specific promoter regions at a crucial time window. In addition, *JMJ706* may play an important role in regulating plant architecture, including plant height. The natural variation in *JMJ706* alleles shows high frequencies across major rice subpopulations, suggesting that *JMJ706* could play an important role in the geographical distribution and adaptation of rice cultivars. Our results may add a new layer to the rice flowering-time regulatory pathway, supporting regional adaptation and potential for future breeding.

## Introduction

Photoperiod is a key factor for regulating plant growth and development. The sensitivity to the number of hours of light and darkness is referred to as the photoperiod sensitivity (PS) and it affects plants’ transition from the vegetative to the reproductive phase. Based on the floral induction as responsive to PS, plants can be classified as short-day (SD) plants, long-day (LD) plants and day-neutral plants (Thomas and Vince-Prue, 1996). Rice (*Oryza sativa L.*), an important field crop since it is the staple food source for half of the world’s population. The rice can be recognized as the model SD plant, which exhibits a strong PS. Its flowering time is accelerated during the SD conditions and delayed under LD conditions, promoting vegetative growth for robust establishment. The adaptation of various degrees of PS allows rice cultivars to distribute and expand cultivation regions through the process of artificial selection. Generally, the regions of rice cultivation are expanded between 53°N to 40°S. The rice cultivars with lower PS responses are characterized by early flowering and are adapted to grow in higher latitudes of temperate regions (Wu et al. 2013). In contrast, cultivars with severe PS responses have been distributed for wide cultivation regions due to later flowering, accompanied with high yield expectations (Xue et al. 2008; Wei et al. 2010; Brambilla and Fornara, 2013).

Rice PS is regulated by the genetic regulation of light and circadian clock receptors. Phytochrome genes act as the main light receptors and the central unit of photoperiod signal transduction to downstream genes (Takano et al. 2005; Takano et al. 2009; Osugi et al. 2011). Therefore, phytochromes play an important role in PS in rice since the phytochrome structure and function-related mutants are highly insensitive to the photoperiod. The null mutants of the *PHOTOPERIOD SENSITIVITY5* (*SE5*) have been reported as a photoperiod-insensitive mutant and the gene encodes the heme oxygenase enzyme involved in phytochrome chromophore biosynthesis (Izawa et al. 2000; Andrés et al. 2009). The *PHOTOSENSITIVITY 13* (*SE13/OsHY2*) encodes phytochromobilin (PΦB) synthase, a crucial enzyme in the phytochrome-chromophore biosynthesis. The null mutants of *SE13* show early flowering and complete insensitivity to the photoperiod (Yoshitake et al. 2015). A long day specific floral repressor *Grain number, Plant height and Heading-date7* (*Ghd7*) which encodes CO, CO-like and TOC1 (CCT)-domain protein positioned downstream of the phytochromes (Xue et al. 2008; Itoh et al. 2010; Osugi et al. 2011). It directly represses a B-type response regulator *Early heading-date 1* (*Ehd1*) that acts as a floral promoter under LD and SD conditions (Doi et al. 2004; Zhao et al. 2015a; Nagalla et al. 2021). The tandem-duplicated, florigen coding *Heading date 3* (*Hd3a*) and *RICE FLOWERING LOCUS T 1* (*RFT1*) genes (Kojima et al. 2002; Komiya et al. 2009) are under the regulation of *Ehd1*. These genes are considered as the orthologs of the *Arabidopsis* florigen gene *FLOWERING LOCUS* T (*FT*) (Corbesier et al. 2007). In addition to the photoperiod signals, the circadian clock signals are crucial for PS response. Rice *GIGANTEA* (*OsGI*) acts as an upstream receptor for circadian clock signals and promotes flowering under both SD and LD conditions (Hayama et al. 2002). The *Heading date 1* (*Hd1*), the rice homolog of *Arabidopsis CONSTANS* (*CO*) (Yano et al. 2000) obtains circadian clock signals from *OsGI* and acts as an important regulator for PS. The rice *Hd1* has been proven to have a photoperiod-dependent function over flowering time control. It generally induces flowering under SD conditions by direct induction of florigen coding genes (Ishikawa et al. 2011). Under LD conditions, *Hd1* participates in a post-translational regulatory complex with *Ghd7* to repress flowering via repressing the *Ehd1* (Nemoto et al. 2016). In addition, CCAAT-box-binding transcription factor *Days-to-heading on chromosome 8* (*DTH8*) has been identified as an LD-specific *Ehd1* repressor (Wei et al. 2010). Previous research has shown that the *Hd1*, *Ghd7*, *DTH8* and their allele combinations play important roles in strong PS. The natural variations of these alleles result in various PS levels and they provide a basis for adaptation of rice cultivars to broad regions (Zong et al. 2021). The *Pseudo-response regulator 37* (*OsPRR37*/*DTH7*) a CCT-domain protein, has been characterized as another *Ehd1* repressor in LD, which ultimately modifies the PS in rice (Gao et al. 2014). The PS directly affects the flowering time decision of the rice plant and thus, influences the plant’s vegetative to reproductive phase transition. Identifying novel genes and deciphering the molecular genetic mechanisms underlying PS would be a crucial goal for rice breeders and plant biologists.

Histone modifications play an important role in gene regulation, including facilitating chromatin unwinding (Pfluger and Wagner, 2007) and recruitment of transcription factors (Field and Adelman, 2020). These major posttranslational modifications are mainly achieved through methylation, acetylation, phosphorylation, and ubiquitylation in both plants and animals (Le et al. 2025). The regulation of histone methylation relies on the coordinated activity of histone methyltransferases, responsible for adding methyl groups, and histone demethylases, which remove them (Williams and Gehring, 2020). Histone methylation is commonly associated with gene silencing in plants. The most abundant histone modifications are histone H3 methylation at lysine 27 (H3K27me) and histone H3 methylation at lysine 9 (H3K9me) (Liu et al. 2010). The outcome of these methylations would be contraction of DNA–histone interactions, followed by limited chromatin access for transcription factors. Methylation at histone H3 lysine 9 mainly takes the forms of H3K9me1 and H3K9me2 (Jacob et al. 2014). These are the positions mainly targeted by the H3K9 methylation erasers known as JUMONJI DOMAIN-CONTAINING PROTEIN, which possess a conserved domain known as Jumonji C (JmjC) (Sun and Zhou, 2008). Previous research has identified a total of 20 JmjC domain containing proteins in rice (Lu et al. 2008) and they were reported to be involved in crucial biological functions. Briefly, JMJ720 was found to be involved in enhancing drought tolerance (Wang et al. 2025b) and a negative regulator of grain size in rice (Wang et al. 2025a). JMJ703 is acting as an important regulator in transposable elements silencing (Cui et al. 2013). The mutants of *JMJ704* were susceptible to *Xanthomonas oryzae pv. oryzae* infection compared to the wild type, indicating JMJ704 induces *Xanthomonas oryzae* resistance (Hou et al. 2015). H3K27me3 demethylase coding JMJ705 interacts with WUSCHEL-RELATED HOMEOBOX 11 (WOX11) for activation of shoot development-related genes in rice (Cheng et al. 2018). Importantly, JmjC domain coding genes have previously been reported on flowering time regulation in rice. The *JMJ701* code for H3K4me3 demethylase that suppresses *RFT1* and delays flowering under natural daylength (ND) conditions (Yokoo et al. 2014). Mutants of the *JMJ706* resulted in a deformed floral organ mainly due to repression of *Degenerated Hell1*(*DH1*). The *JMJ706* code for the H3K9me2 and H3K9me3 demethylase, which induces an activator effect on the target gene (Sun and Zhou, 2008). In addition, a recent report explains that *JMJ706* functions as a flowering time regulator and mutants improve grain yield by extending regional adaptability (Yu et al. 2025).

In the current study, we investigated the PS regulatory function of the *JMJ706* gene in detail. It affected rice photoperiodic flowering time regulatory network in a day length-dependent manner through promoting *Ghd7* expression under LD and suppressing *Ehd1* expression under SD conditions. The impact of the H3K9me2 demethylation activity may extend to plant architecture and grain yield-related traits regulation. Moreover, the natural variations of the *JMJ706* alleles tell us that the specifically dominant alleles facilitate the regional adaptability of rice varieties. Therefore, our findings may add a novel PS pathway through epigenetic regulation of key flowering time genes in rice.

## Results

### *JMJ706* is a candidate photoperiod-sensitive gene

The carbon-ion beam induced mutant 13C3-97 shows relatively early flowering compared to Nipponbare (WT) plants in field conditions. To examine the flowering time behavior in detail, we grew the 13C3-97 mutant in artificial SD and LD conditions. Under SD conditions, the 13C-97 showed an approximately 18-day delay, whereas under LD conditions, flowering was accelerated by 14 days relative to WT plants. Based on the flowering time behavior under SD and LD conditions, the photoperiod sensitivity indexes (PSI) are 0.19 and 0.45 in 13C3-97 and WT plants, respectively (Supplementary Fig. S1A). The low PSI suggested the candidate mutated gene in 13C3-97 could be a photoperiod-sensitivity gene. We performed whole-genome sequencing analysis of the 13C3-97 mutant, and four candidate causal genes were identified (Supplementary Table S1). Among them, *Os10g0577600* (*JMJ706*/*Jumonji 706*), which carries a homozygous 1-bp deletion and encodes an H3K9 demethylase, was considered a strong candidate based on findings from previous reports showing that Jumonji family genes play roles in flowering time control and photoperiod sensitivity in rice (Yokoo et al. 2014). To identify the causal gene of the mutant, we created CRISPR-Cas9 knockout lines for *JMJ706*. Among the knockout lines, we selected jmj-5 and jmj-6, which possess a 14-bp deletion and 1-bp insertion, respectively, in the JmjC domain coding region (Supplementary Fig. S1B, Fig. 1B). In addition, we grew a BC_1_F_2_ population derived from a cross between the 13C3-97 and WT plants. A progeny carrying a homozygous mutation in the *JMJ706* from the F_2_ population was selected and named jmj-M. These plants possessed a 1 bp deletion in *JMJ706*, which leads to the loss of C5HC2 Zinc-finger domain (Supplementary Fig. S1B, Fig. 1B). A genetic linkage analysis using heading date as a phenotypic trait was conducted under natural-day-length conditions. The early-flowering phenotype was linked to *JMJ706*, suggesting that *JMJ706* may be responsible for the early-flowering phenotype. These jmj-M, and jmj-5 and jmj-6 (T_2_ generation) plants were grown under SD and LD conditions and flowering time was observed. Consistently, a clear acceleration of flowering under LD conditions and a delay of flowering under SD conditions, compared to WT was observed in all three mutant lines (Fig. 1 A, C and D). Their flowering time difference under SD and LD conditions was not significant, leading to lower PSI values (Fig. 1A). However, PSI of 13C3-97 is greater than these mutants, indicating the 13C3-97 may possess other mutations that possibly influence flowering time regulation. Based on these results, the *JMJ706* gene can be considered a photoperiod sensitivity regulatory gene in rice. Since the growth rate influences the flowering time regulation, we measured the leaf age of both mutants under SD and LD conditions. Both jmj-5 and jmj-6 mutants resulted a significantly lower leaf age, indicating a slower growth rate compared to WT under both conditions (Fig. 1E).

**Figure 1.**
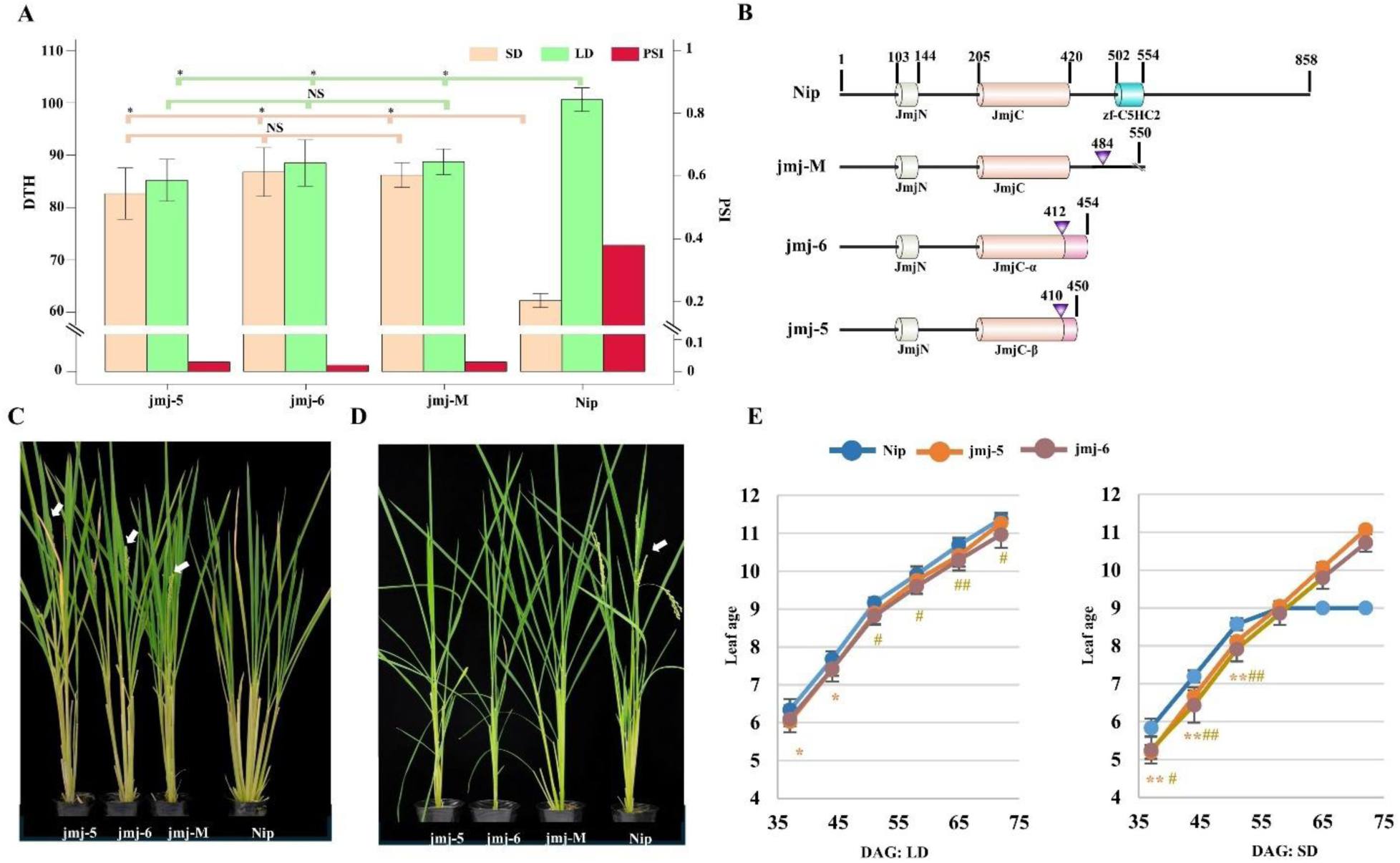
*JMJ706* regulates photoperiod sensitivity and plant growth rate. **A)** Days to heading (DTH) and photoperiod sensitivity index (PSI) of CRISPR-Cas9 mutants (jmj-5 and jmj-6), Carbon-ion induced mutant (jmj-M) and Nip plants under SD and LD conditions (\**P* < 0.01 and the NS stands for “Not Significant”). **B)** Graphical representation of the amino acid (AA) sequence alignment and protein domain annotation of the JMJ706 protein of each mutant and Nip. Numbers indicate the number of AA and the inverted triangle indicates the position of the mutation. The extended mutated AA sequence is indicated in a different color and \\ indicates the position of the AA sequence termination. The symbols of α and β indicate the mutated forms of each JmjC domain possessed by jmj-5 and jmj-6 mutants, respectively. **C)** *JMJ706* mutants and Nip plants under LD conditions. Photos were taken when the mutants flowered around 88-days after sowing. **D)** *JMJ706* mutants and Nip plants under SD conditions. Photos were taken when the Nip flowered around 85-days after sowing. White color arrows indicate the immature panicles. **E)** Leaf-age measurements of *JMJ706* mutants and Nip plants under LD and SD conditions. Data are means ± S. D. (n > 3 biological replicates), and the significance of the difference was assessed by Student’s t-test (** and ## *P* < 0.01; * and # *P* < 0.05. The symbols * and ** stand for jmj-5 and # and ## stand for jmj-6).

### *JMJ706* is a floral repressor under LD conditions and a floral promoter under SD conditions

To elucidate the *JMJ706*’s role in regulating photoperiodic flowering, the relative gene expression of key flowering time and circadian clock genes was observed 2 hr after dawn, under both LD and SD conditions. As expected, the *Ghd7* expression was significantly reduced in both jmj-5 and jmj-6 mutants under LD conditions (Fig. 2). The reduced *Ghd7* expression triggers the induction of *Ehd1* and thus subsequently induces *Hd3a* and *RFT1*. This induction of florigen coding genes in *JMJ706* mutants compared to WT under LD conditions could be the reason for the early flowering phenotype. In contrast, under SD conditions, *Ehd1*, *Hd3a* and *RFT1* resulted in a significant expression reduction in jmj-5 and jmj-6 compared to WT, indicating the possibility of low florigen production leading to later flowering. Altogether, these spot gene expression data are consistent with the photoperiod-insensitive phenotype of *JMJ706* mutants.

**Figure 2.**
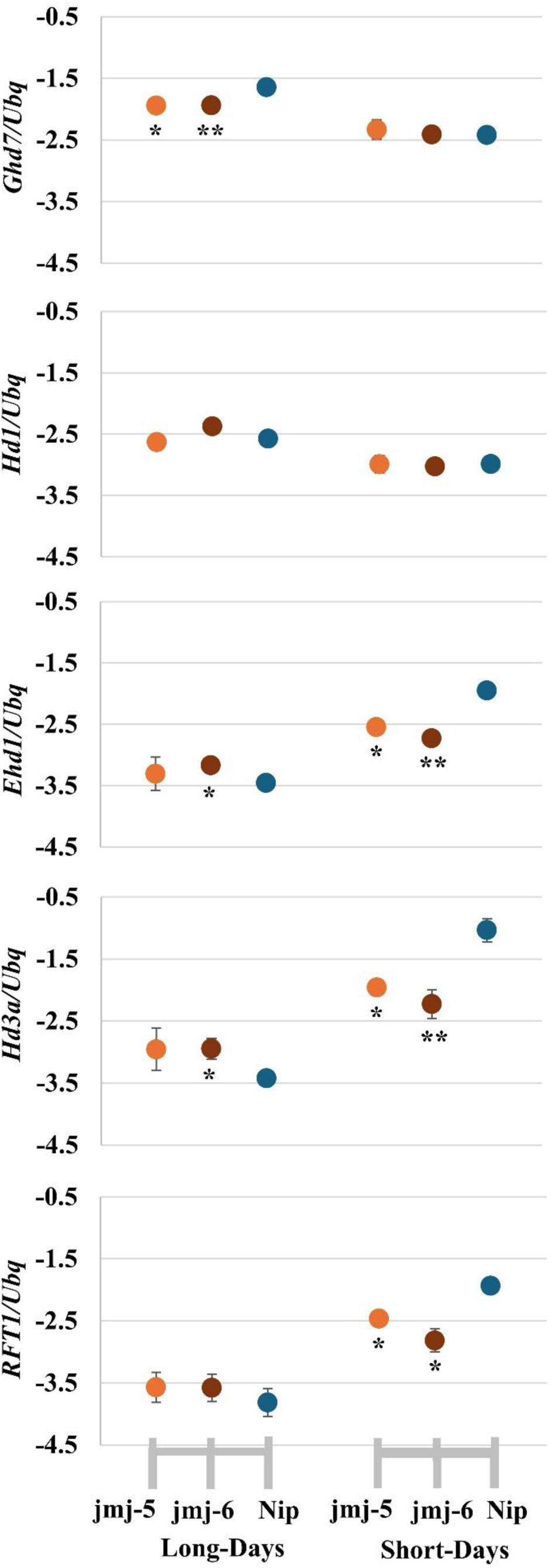
Spot relative gene expression of *Ghd7*, *Hd1*, *Ehd1*, *Hd3a* and *RFT1* in *JMJ706* mutants and Nipponbare plants. Leaf blades of three weeks old seedlings were harvested 2hr after dawn. Data are means ± S. D. (n = 3 or 4 biological replicates), and the significance of the difference was assessed by Student’s t-test (\*\**P* < 0.01; \**P* < 0.05). Relative gene expression was shown in the logarithmic Y-axis.

Since *JMJ706* has a significant impact on the key flowering time regulatory genes, a detailed investigation of its expression dynamics was performed using a diurnal gene expression analysis. Under both SD and LD conditions, leaf blade samples of WT and jmj-5, targeting eight sampling points within 24 hr, were collected (Fig. 3O). The *Ghd7* expression resulted in a significant reduction throughout the LD day-time in jmj-5. As a response, the *Ehd1* expression has peaks in the early morning and at midnight. Since The *Ehd1*, act as a floral promoter under LD and SD conditions, the downstream *Hd3a* and *RFT1* show a clear induction around early morning (Fig. 3: A, C, E and G). Both diurnal gene expression analysis (Fig. 3) and the spot gene expression analysis (Fig. 2) pointed out that *Ghd7* mRNA reduction under LD conditions could be the key factor that triggers the induction of florigen coding genes in jmj-5 mutants. Therefore, under LD conditions, the functional *JMJ706* could be act as a floral repressor mainly by promoting *Ghd7* expression. Under SD conditions, jmj-5 exhibit a significant *Ehd1* reduction around mid-night and early morning compared to WT. Since the *Ehd1* positively regulates the *Hd3a* and *RFT1*, both genes were repressed and resulted a later flowering phenotype (Fig. 3: D, F, and H). The functional *JMJ706* mainly promotes the *Ehd1* expression under SD mid-night conditions and functions as a candidate floral promoter.

**Figure 3.**
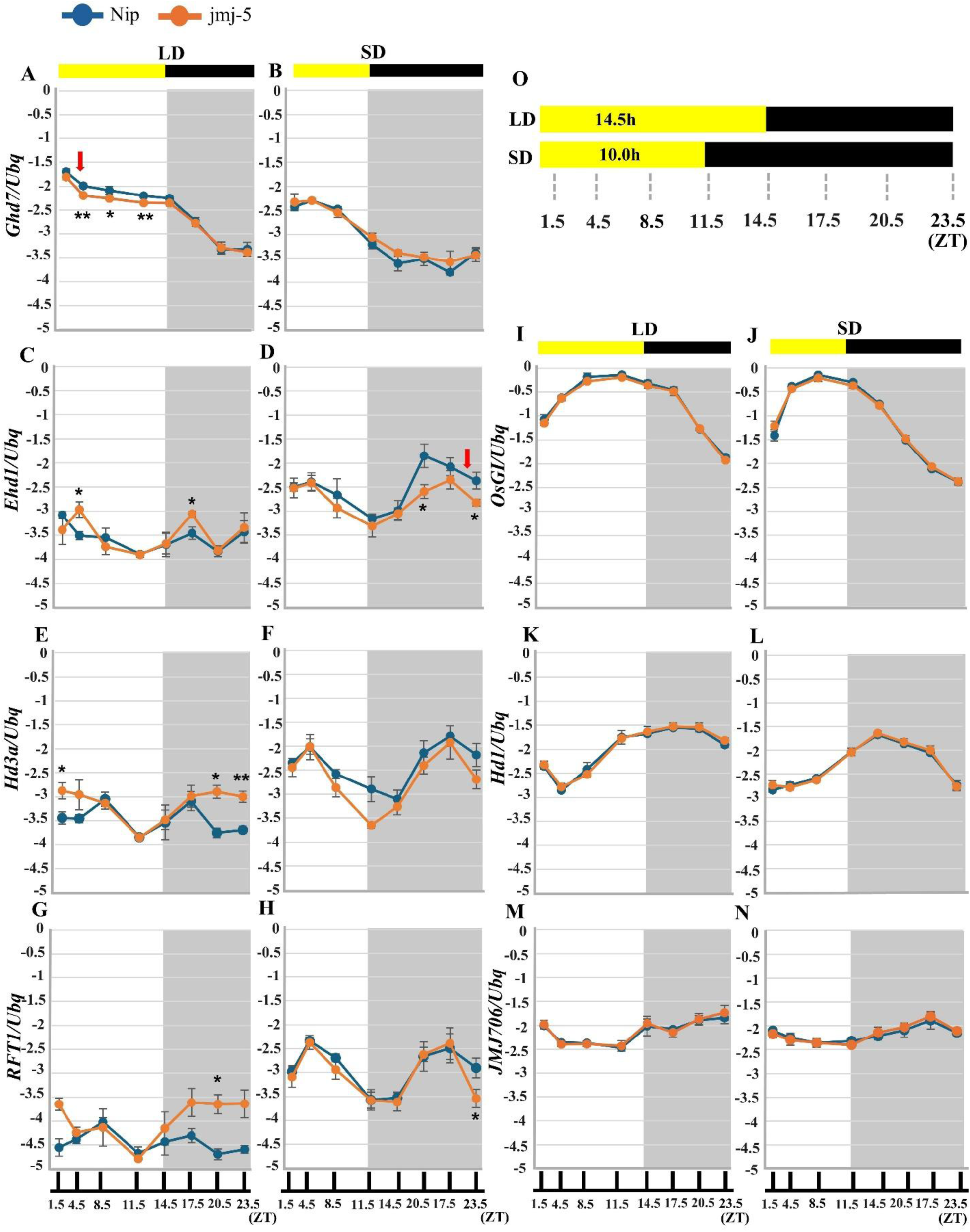
Diurnal relative gene expression of *Ghd7*, *Ehd1*, *Hd3a, RFT1*, *OsGI*, *Hd1* and *JMJ706*. Plant materials are three-week-old seedlings of jmj-5 and Nipponbare grown under LD **(A, C, E, G, I, K, M)** and SD **(B, D, F, H, J, L, N)** conditions. **O)** Yellow and black horizontal bars represent light and dark periods, respectively. Data are means ± S. D. (n = 3 or 4 biological replicates); and the significance of the difference was assessed by Student’s t-test (\*\**P* < 0.01; \**P* < 0.05). Relative gene expression was shown in the logarithmic Y-axis.

The *Hd1* is positioned as an upstream gene of *Ehd1* and involved in modifying its expression under SD and LD conditions (Ishikawa et al. 2011; Nemoto et al. 2016). Therefore, we checked the diurnal expression of *Hd1* to identify the impact of the *JMJ706* mutation on its expression. There is no significant gene expression difference of *Hd1* in jmj-5 compared to WT during the 24 hr, indicating *JMJ706* does not modify its transcription (Fig. 3: K and L). Interestingly, *OsGI* diurnal gene expression is not altered by the *JMJ706* mutation, demonstrating *JMJ706* may function downstream of *OsGI* (Fig. 3: I and J). The diurnal gene expression of *JMJ706* follows a circadian rhythm, suggesting its function as a circadian clock responsive gene. Additionally, the native expression dynamics of *JMJ706* indicated that its transcript levels are induced in the dark phase and reduced during the light irrespective of the lay length conditions (Fig. 3: M and N).

### *JMJ706* facilitate *Ghd7* and *Ehd1* expression through H3K9me2 demethylation on the promoter region

The diurnal gene expression analysis strongly proposed that *JMJ706* conveys photoperiod sensory signals to the flowering time regulatory network via *Ghd7* and *Ehd1*. Since *JMJ706* coding for H3K9me2 demethylase and proven to facilitate target gene expression (Sun and Zhou, 2008), we hypothesize that H3K9me2 demethylase activity may trigger those gene expressions based on circadian clock signals. Specifically, *Ghd7* and *Ehd1* diurnal gene expression analysis revealed candidate critical time points when the demethylase activity may exclusively promote their gene expression. Under LD conditions, a significant difference of *Ghd7* expression between WT and jmj-5 resulted around 4.5 hr after dawn. We collected leaf blade samples at 4 hr after dawn (Fig. 3: A (red arrow). Under SD conditions, a significant difference of *Ehd1* expression was 0.5 hr before dawn and leaf sample collection was conducted 1 hr before dawn (Fig. 3: D (red arrow).

In the jmj-5 mutant, defective H3K9me2 demethylase may fail to perform the demethylation activity and H3 proteins remain tightly bound to the promoters of the target genes. Therefore, the anti-H3K9me2 antibody should precipitate a higher amount of DNA in a specific region on the promoter (or genic) compared to WT. To assess this possibility, a ChIP-qPCR experiment was performed using LD leaf samples and searched for candidate positions on the *Ghd7* gene region. A significantly higher amount of DNA has precipitated in jmj-5 compared to WT on the 291 bp and 655 bp upstream of the transcription start site (TSS) (Fig. 4A). The ChIP-qPCR experiment was extended using leaf samples from SD conditions and the *Ehd1* gene region was targeted. A significantly higher DNA precipitation was observed near the TSS: 107 bp downstream, in jmj-5 compared to WT (Fig. 4B). These ChIP-qPCR results suggest that *JMJ706* enhances *Ghd7* and *Ehd1* expression through demethylation of specific regions of their promoter.

**Figure 4.**
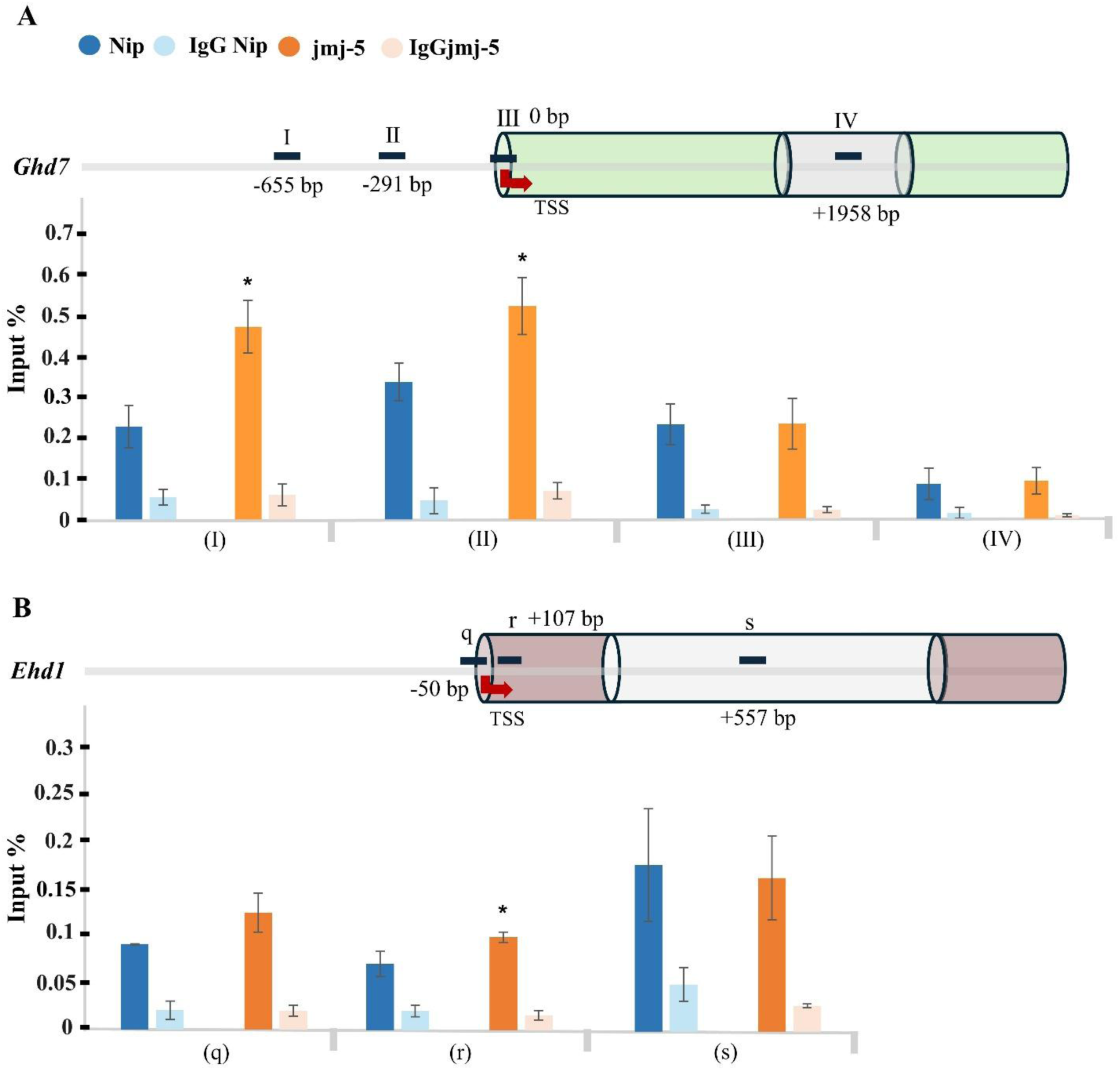
JMJ706 facilitates chromatin relaxation around *Ghd7* **(A** under LD) and *Ehd1* **(B** under SD) promoter regions. The figure shows the data from chromatin immunoprecipitation followed by quantitative PCR (ChIP-qPCR). The upper section of each A and B shows the graphical representation of the fraction of the genomic region: exons (thick color) and intron (light color) of *Ghd7* and *Ehd1,* respectively. Black horizontal lines indicate the relative positions of PCR amplicons. Red arrows indicate the transcriptional start site (TSS). IgG antibody was used as a negative control (mock). The quantities of immunoprecipitated DNA were normalized to those of the input DNA and the y-axis label is % of input DNA. Data is from at least three biological replicates and means ± S. D. The significance of the difference was assessed by Student’s t-test (\**P* < 0.05).

### *JMJ706* regulates multiple growth and development related genes

Since *JMJ706* encodes one of the key epigenetic regulatory enzymes, we are interested in investigating other candidate target genes regulated by *JMJ706*. This assumption was further supported by the observation of significantly higher plant height in *JMJ706* mutants under LD conditions compared to WT plants (Fig. 5E). However, under SD conditions, only the jmj-M mutant resulted a significantly higher plant height (Supplementary Fig. 2). In addition, we observed the defective floral organs in all the *JMJ706* mutants as initially described by the Sun and Zhou, (2008) (Supplementary Fig. 3). Diurnal gene expression analysis resulted a day-length dependent, circadian clock responsive gene expression dynamic of *JMJ706* (Fig. 3: M and N). Therefore, we collected the total RNA at 0.5 hr and 3.5 hr before dawn under LD and SD conditions, respectively, targeting the interval around the *JMJ706* peak gene expression. First, we searched for possible flowering time and circadian clock regulatory genes within these differentially expressed (DE) genes (Fig. 5 A, B, and C), as a confirmation of previous gene expression analysis. As expected, floral promoters such as *Ehd1*, *Hd3a* and *MADS-box* 14 (*OsMADS14*) were significantly downregulated and *LEC2/FUSCA3-like 1* (*OsLFL1*), a floral repressor, was significantly upregulated in jmj-5 compared to WT under SD conditions (Table 1). However, under LD conditions, we could not find any significaly DE genes that were reported to have a direct affiliation with flowering time or circadian clock.

**Figure 5.**
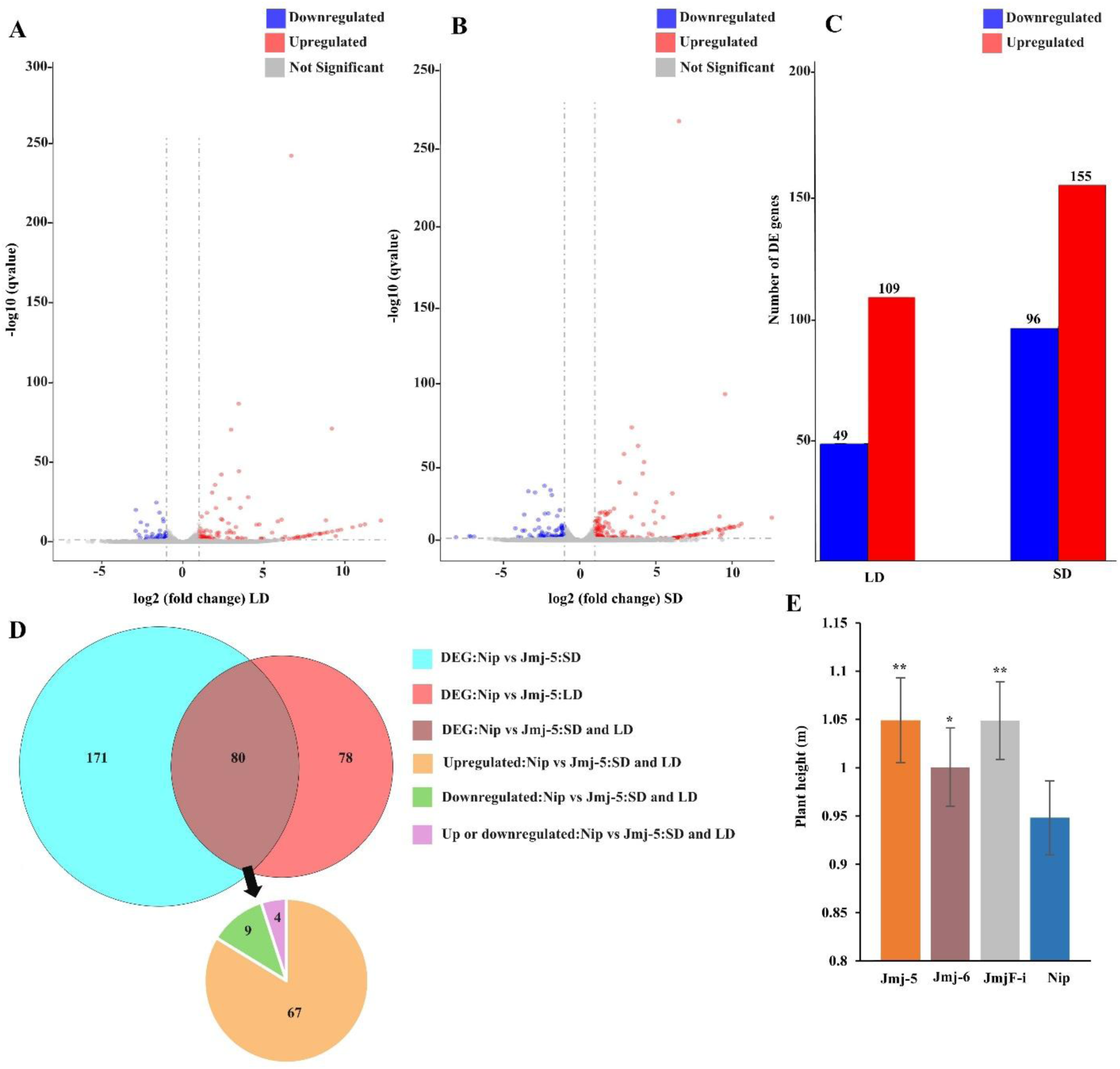
The impact of the *JMJ706* mutation on the mutant transcriptome and rice plant height compared to Nipponbare (Nip). The volcano plot shows the distribution of significantly up and downregulated differentially expressed genes (DEGs) under **(A)** LD and **(B)** SD conditions. **C)** The number of up and downregulated DEGs in the mutant compared to Nip under each SD and LD. **D)** Composition of the up and down-regulated DEGs based on the daylength conditions. **E)** jmj mutants show higher plant height compared to Nip under LD conditions. Data are means ± S. D. (n>3), and the significance of the difference was assessed by Student’s t-test (**P < 0.01; *P < 0.05).

**Table 1:**
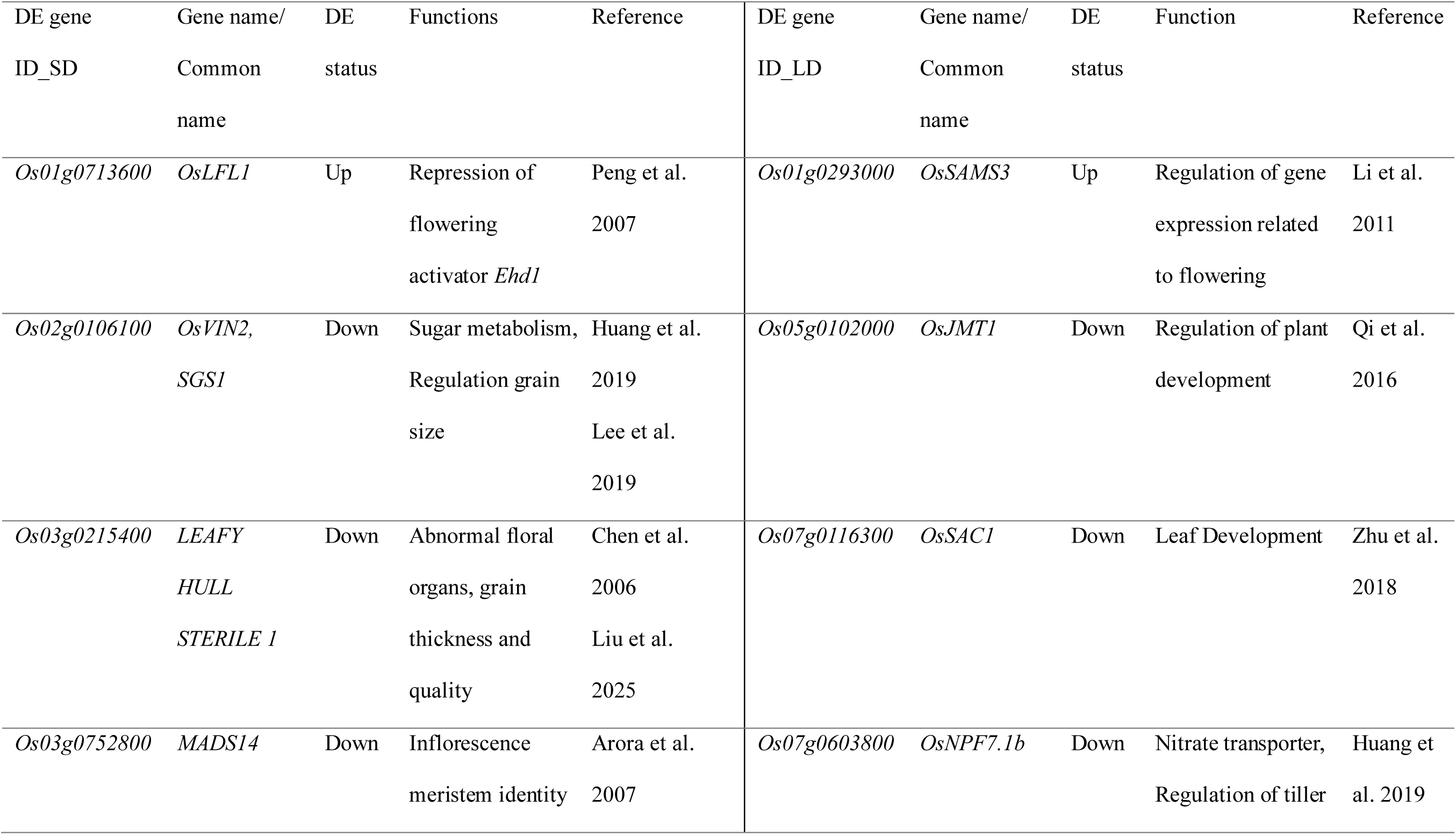
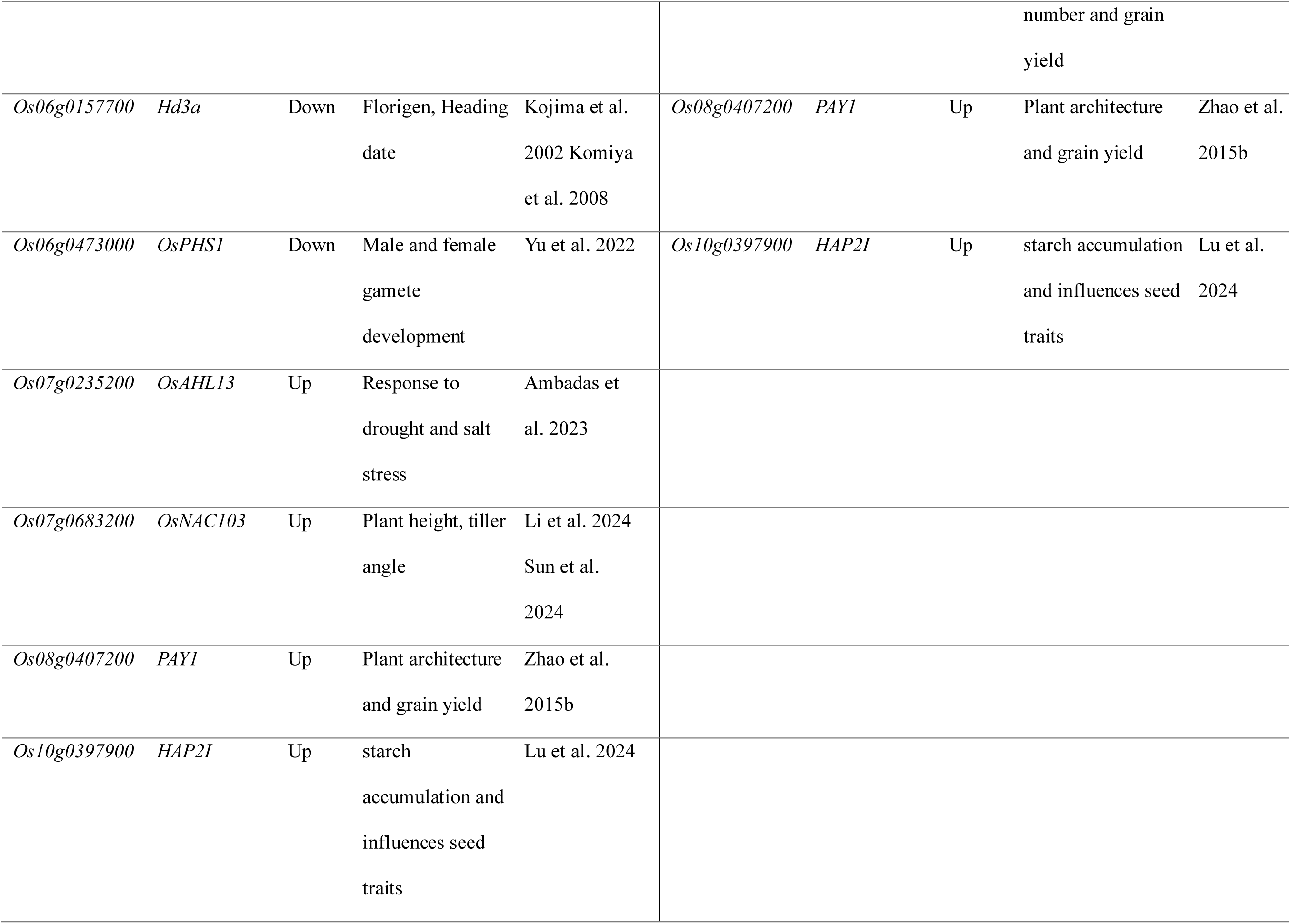
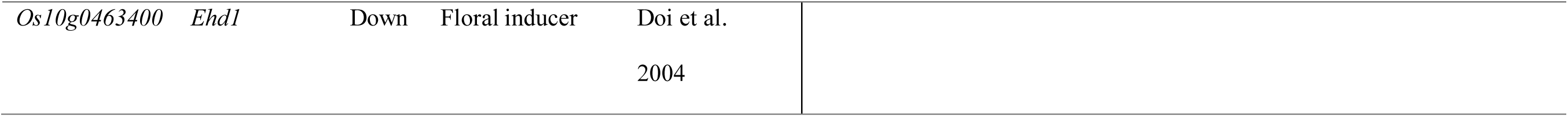
Selected Differentially expressed (DE) genes under SD and LD conditions in Nip vs Jmj5 and their predicted functions in rice.

In addition to the flowering time regulation, important DE genes responsible for seed/grain properties and plant architecture regulation were identified. The *LEAFY HULL STERILE 1*, *PLANT ARCHITECTURE AND YIELD* 1 (*PAY1*), Nuclear factor Y (NF-Y) transcription factor A8 (*HAP2I*) and Vacuolar acid invertase (*OsVIN2*) genes were previously reported for their functions on rice seed-related traits regulation. Candidate genes involved in plant architecture regulation were *PAY1*, *OsNAC103*, *OsSAC1* and *OsJMT1*. In addition, *OsPHS1* and *LEAFY HULL STERILE 1* were annotated for their functions in floral organ development (Table 1). A total of 80 genes were identified as commonly DE under SD and LD conditions, while 67 of them were upregulated and nine were downregulated (Fig. 5D). The *PAY1* and *HAP2I* genes were significantly upregulated under both SD and LD conditions, suggesting *JMJ706* may affect seed and plant architecture-related genes regardless of daylength conditions (Table 1).

### Natural variations of *JMJ706* alleles

The transcriptome analysis discovered the clues that support the considerable impact of the *JMJ706* on controlling multiple key physiological traits in rice. Therefore, we are interested in exploring the natural variations of *JMJ706* alleles across the main rice populations: Temperate Japonica (TEJ), Tropical Japonica (TRJ), Japonica (JP; not distinguished as either TEJ or TRJ), Indica (IND), Aus (AUS) and Aromatic (ARO). The DNA sequences of the transcript region of 409 rice accessions were used to identify 15 single nucleotide variants and three deletions. These variations produce six unique allele types (haplotype groups) (see the method section). Among these variations, there were 11 variations located in the coding region and nine of which were nonsynonymous. Only the haplotype Ⅵ possesses a synonymous codon variant in the JmjC domain. The haplotype Ⅳ, Ⅴ and Ⅵ contained nonsynonymous codon variants in the zinc-finger domain. Interestingly, we could not find a haplotype group that contained a nonfunctional allele of *JMJ706* (Fig. 6A). The haplotype network analysis resulted in two distinguished haplotype clusters. A Cluster with the haplotype groups Ⅴ and Ⅵ is mainly dominated by the individual accessions belonging to the IND sub-population. The other cluster mainly consists of Japonica sub-population accessions. It contained two major haplotype groups: haplotype Ⅰ, dominated by TEJ and haplotype Ⅳ, dominated by TRJ accessions (Fig. 6B and 6C). Generally, *JMJ706* haplotype clusters and individual haplotype groups are clearly dominated by the accessions from a major sub-population.

**Figure 6.**
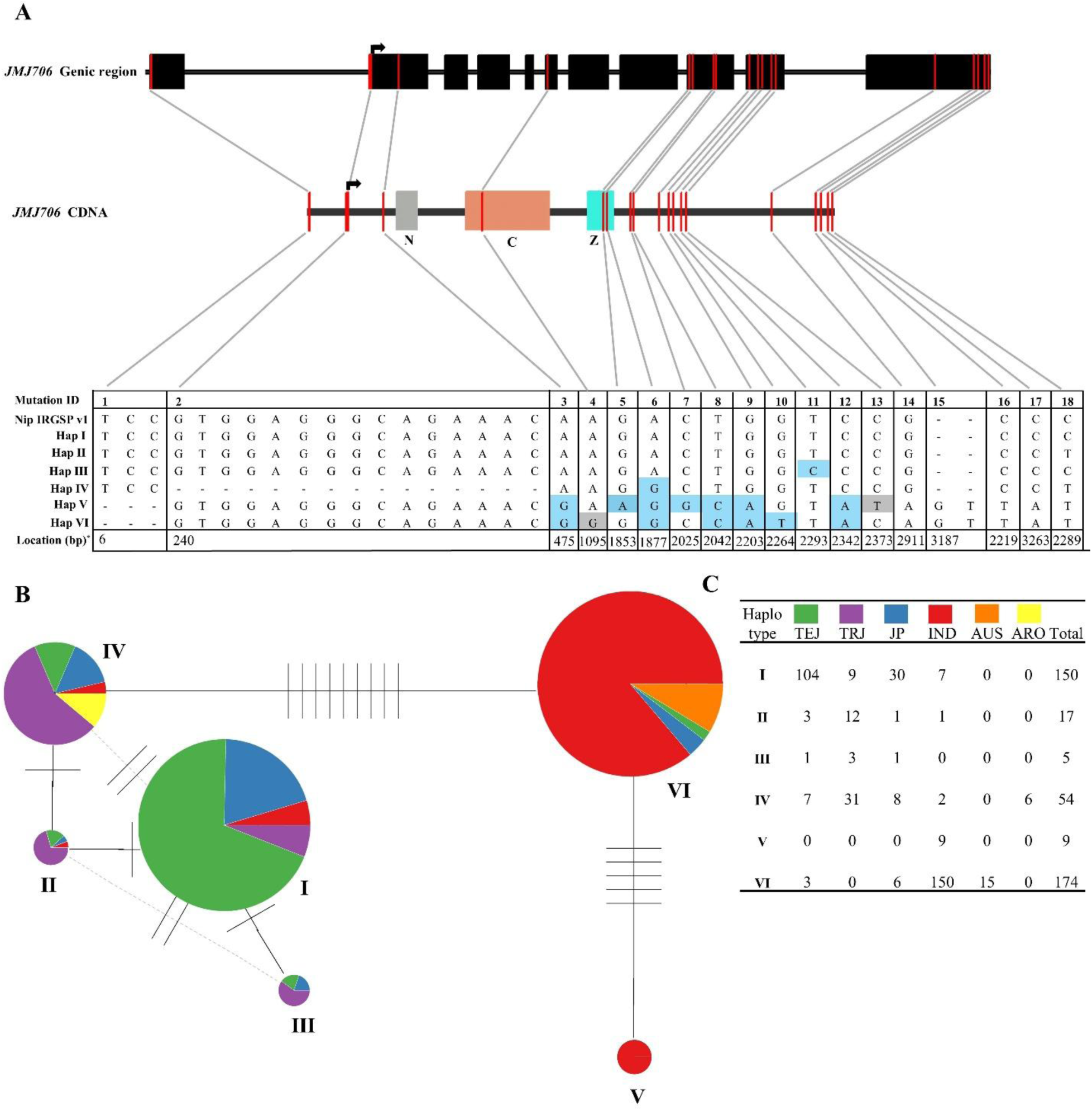
Identification of the main haplotypes of *JMJ706* and their distribution across the major rice sub-populations. A) The top layer shows a graphical representation of the *JMJ706* genic region and black boxes represent the exons. The middle layer represents the complementary DNA (cDNA) of the *JMJ706*. The boxes named N, C and Z represent the regions responsible for coding JmjN, JmjC and the zinc-finger domain, respectively. The red lines indicate the positions of the mutations (single-nucleotide variations and deletions) and the black arrow indicates the position for the translation start site. Bottom layer shows a table that summarizes the haplotype groups, the nature of responsible mutations and the location estimated with respect to the transcription start site. The gray color lines represent and connect the candidate locations for each mutation among the upper two layers. The bases highlighted with light blue and gray represent the nonsynonymous and synonymous mutations, respectively. **B)** Haplotype network which generated based on the identified haplotypes using 409 rice varieties. Each pie chart represents a haplotype group consisting of major sub-populations. The number of lines on the straight line connecting each pair of circles represents the number of single nucleotide variations. The size of the circles represents the number of accessions included in each haplotype group. **C)** Composition of each haplotype group, including the number of accessions from major rice subpopulations.

## Discussion

A comparative genome analysis of JmjC domain-containing proteins among humans, Arabidopsis, and rice were separated into seven major groups. Withing these groups, five sub-groups were identified based on the abundance of the plant proteins. The JMJ706 was grouped to JMJD2/JHDM3 subgroup (Sun and Zhou, 2008), where the members actively reverse lysine methylation, H3K36 and/or H3K9 as their histone substrates. A total of five rice proteins were isolated that featured with JmjN, JmjC and a single C5HC2 (zinc-finger) domain (Fig. 1B) (Lu et al. 2008; Sun and Zhou, 2008; Chen et al. 2011). In this study, two CRISPR-Cas9 mutants generate a mutated, truncated JmjC domain and completely abolish the zinc-finger domain. The jmj-M mutant has an intact JmjC domain but it also loses the zinc-finger domain. However, all these mutants displayed a clear and comparable photoperiod insensitivity, indicating that the importance of the zinc-finger domain might be greater than JmjC domain regarding the PS regulation. A recent research produced a mutant called *ispl10*, the causal gene is the *JMJ706*, which possessed a truncated ISPL10 protein that lacked the entire zinc-finger domain. These mutants showed later flowering under SD and early flowering under LD conditions, indicating Photoperiod insensitivity (Yu et al. 2025). Further evidence was provided by the JMJ703/JMJ704 binding with the *OsLFL1* promoter. The H3K4me3 demethylation and the promotion of H3K27me3 levels at *OsLFL1* chromatin were eliminated when deleting the zinc-finger domain (Xuan et al. 2024). During the initial identification of *JMJ706*, Sun and Zhou (2008) used T-DNA-inserted knockout lines, which eliminate JMJ706. These mutants displayed characteristic floral organ defects and all three mutants, jmj-M, jmj-5 and jmj-6, generated in this study were also shown to have comparable floral organ defects (Supplementary Fig. 3). However, the lack of H3K9me2 or H3K9me3 demethylase does not impose a completely defective floral organ containing flowers in a panicle, as we got the filled grains and the germination ratio of the filled grains was satisfied (data not shown). These results indicate that the *JMJ706* mutation induces a partial effect on the floral organ development regulatory genes (*DH1* and *OsMADS47* (Sun and Zhou, 2008)). But it has a serious effect on PS regulatory genes and results in complete photoperiod insensitivity in rice.

Rice heading date regulation is managed mainly through two distinguished pathways. The evolutionary conserved pathway: *OsGI*-*Hd1*-*Hd3a*/*RFT1* and monocot/rice specific pathway: *Ghd7*-*Ehd1*-*Hd3a*/*RFT* (Tsuji et al. 2011). In the current study, we found that *JMJ706* is associated with the rice-specific heading date regulatory pathway to modify PS response (Fig. 7). According to the mode of action of *JMJ706*, it activates two critical genes in distinct time frames. Under LD, the demethylase activity promotes *Ghd7* expression in the morning (Fig. 3A) by relaxing the chromatin around the promoter region (Fig. 4A). Since *Ghd7* is a floral repressor, it represses downstream floral promoters *Ehd1*-*Hd3a/RFT*. Interestingly, no *Ghd7* expression promotion was observed under SD conditions compared to jmj-5 (Fig. 3B). This suggests that *JMJ706*-mediated *Ghd7* promotion occurs exclusively under LD conditions. Moreover, *JMJ706* native mRNA levels do not show a clear difference and do not peak either in LD or SD morning (Fig. 3M and 3N). Therefore, JMJ706 (H3K9me2 demethylase) is expected to undergo day-length-dependent posttranslational modification. In contrast, Yu et al. (2025) suggest that the *JMJ706* facilitated LD flowering does not depend on the *Ghd7* using *ISPL10* mutants. However, in this study, we have verified the fact that *Ghd7* expression can be significantly enhanced by *JMJ706* using independent mutants that appeared in multiple experiments (Fig. 2 and Fig. 3A).

**Figure 7.**
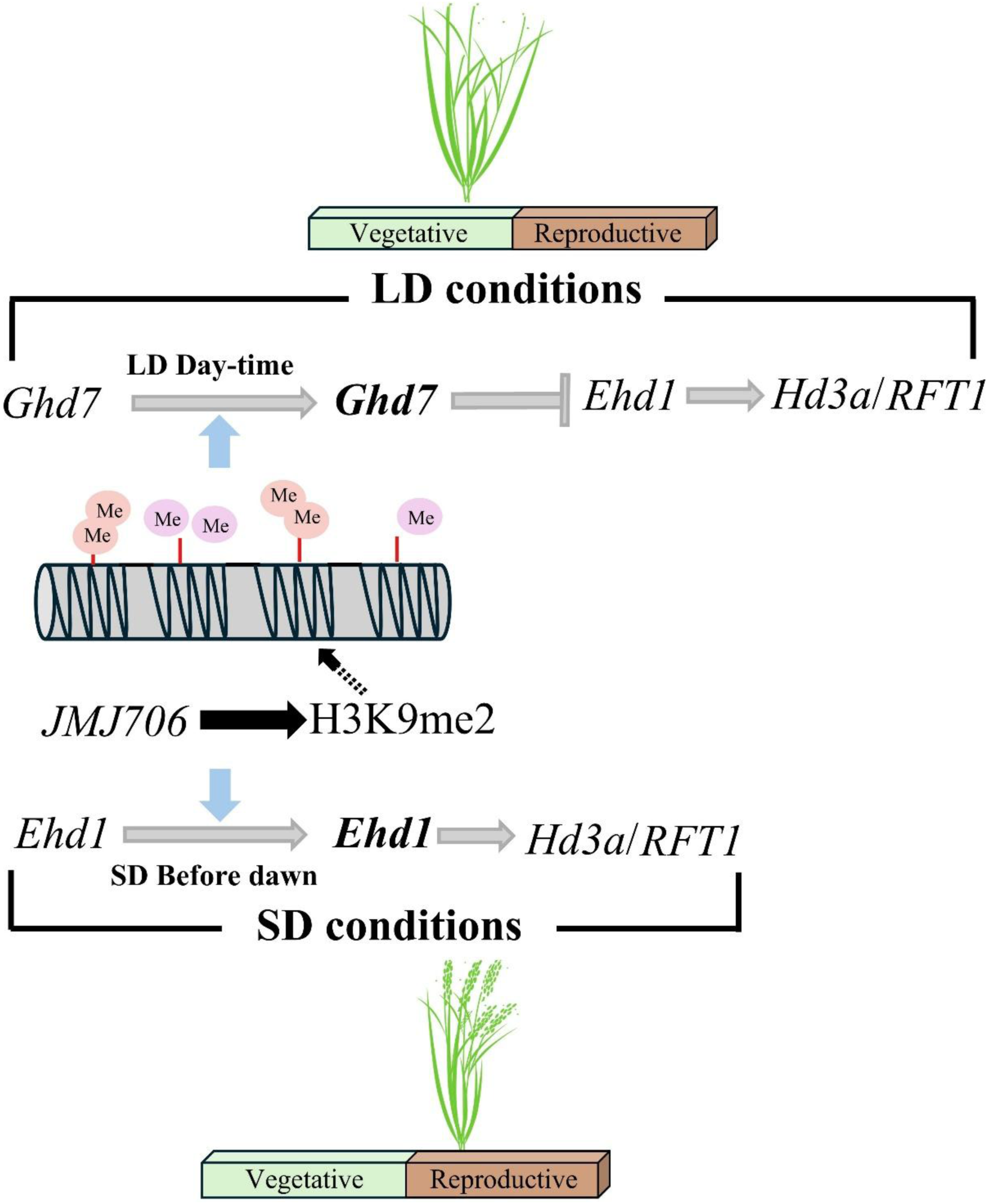
Proposed model of *JMJ706* involvement for LD and SD dependent flowering time regulatory pathway in rice. The gray pointed arrows and flat-head arrows indicate the transcriptional activation and repression, respectively, while the black arrow shows the translation of *JMJ706* to H3K9me2. The highlighted ***Ghd7*** and ***Ehd1*** represent the transcriptionally activated genes due to H3K9me2 demethylase function (represented using blue arrows). The black-dashed arrow indicates post-translational modification on the histone. The light orange color Me and pink color Me represent histone-bound methyl groups and demethylated methyl groups, respectively. Briefly, under LD, *JMJ706*-encoded H3K9me2 promotes *Ghd7* transcription, further repressing *Ehd1* and suppressing *Hd3a* and *RFT1* to facilitate later flowering and maintain vegetative growth. Under SD, H3K9me2 promotes *Ehd1* transcription, induces the florigen coding *Hd3a* and *RFT1* and facilitates early flowering. Therefore, *JMJ706* accelerates the reproductive phase under SD conditions.

Under SD, *Ehd1* appeared to be the direct target for *JMJ706* since its expression is greatly promoted around midnight (Fig. 3D). We found that a significantly higher H3K9me2 demethylase activity near the TSS of *Ehd1* (Fig. 4B). Since *Ehd1* acts as a floral promoter under SD, it promotes *Hd3a/RFT* and stimulates flowering (Fig. 7). However, we could not confirm the *Ehd1* promotion by *JMJ706* under LD conditions due to the excessive repressor effect of *Ghd7* (Fig. 3C). In addition to *Ehd1* and *Hd3a*, transcriptome analysis picked up two candidate causal genes responsible for floral promotion under SD. The *OsLFL1* acts as an *Ehd1* repressor under SD conditions (Peng et al. 2007) and *JMJ706* downregulates *OsLFL1,* derepressing *Ehd1*. The *OsMADS14* acts as a floral promoter (Jeon et al. 2000) and *JMJ706* significantly upregulates its mRNA levels (Table 1). Remarkably, the *JMJ706* does not modify the key gene expression related to evolutionarily conserved pathway: *OsGI* and *Hd1,* based on the solid diurnal expression data (Fig. 3 I,J,K and L). Collectively, we portray the *JMJ706* as an important photoperiod sensor in rice since it stimulates flowering in short photoperiods and depresses flowering under long photoperiods via fine-tuning of heading date regulators.

Our study further extended to investigate additional phenotype features regulated by *JMJ706*. We measured the leaf age in jmj-5, jmj-6 mutants and WT plants under SD and LD conditions. The objective was to assess whether the heading date regulation of *JMJ706* was accomplished through genetic manipulation of related genes or by regulating the plant growth rate. Under SD conditions, WT plants show a higher growth rate, and they flower earlier compared to mutants. In contrast, under LD conditions, WT plants flowered later, but still the plants showed a higher growth rate (Fig. 1E). Based on these results, *JMJ706* indeed manipulates the plant growth rate; however, it does not depend on the day-length conditions. Moreover, *JMJ706*-affiliated heading date regulation does not depend on the growth rate since WT plants were later flowering, yet they show higher growth rates compared to mutants under LD conditions (Fig. 1A and 1E). In addition, our results suggested that *JMJ706* plays an important role in regulating plant architecture. Since all the mutants show higher plant height compared to WT plants, *JMJ706* functions as a repressor for plant height under LD. Although we expected the same phenotype under SD, only the jmj-M showed a significant height compared to WT, probably due to the intact JmjC domain, which might be important in plant height regulation under SD (Fig. 5E and Supplementary Fig S2).

The study of natural variation of *JMJ706* resulted in six unique allele types among the major rice populations. Based on our results, JmjN and JmjC domains are conserved and only the zinc-finger domain possesses a structural mutation in two positions. However, those are nonsynonymous mutations altering a single amino acid modification and do not modify the entire AA sequences. These results imply the importance of the functional *JMJ706* activity across the entire rice population. The domination of rice sub-population within haplotype groups indicates a clear population-specific adaptation. A given sub-population being enriched in a haplotype group suggests that it likely conferred selective advantages under that group’s ecological or agronomic environments (Zhang et al. 2015; Cui et al. 2020). Therefore, haplotype Ⅴ and Ⅵ are mainly dominated by Indica type accessions having a mutation in a zinc-finger domain (mutation ID 5 and 6 in Fig. 6). This mutation may alter the JMJ706 function, lower the PS and be expanded in tropical areas where the day length is relatively comparable throughout the year. It is noteworthy that the mutants produced in this study also contained a deleted zinc finger domain and they show a higher degree of photoperiod insensitivity. In contrast, haplotype Ⅰ, dominated by temperate Japonica accessions, where the PS is crucial for their survival in temperate geographical areas where the daylength difference is clearly observable. Thus, having an intact and functional JMJ706 could be advantageous for proper PS and proper adaptation to these environments. Remarkably, the *JMJ706* null mutants have likely been eliminated from the major rice populations since they carry floral organ defects, and such a haplotype may not be sustainable (Supplementary Fig 3). Based on these evidences, we propose that *JMJ706* may play a key role in the geographic distribution and environmental adaptation of the main rice populations.

## Methods

### Plant materials, growth conditions and measuring days to heading

The rice (*Oryza. sativa* L.) cultivar Nipponbare (Nip) was used as the WT. Dry rice seeds of Nipponbare were irradiated by C-ion beams (^12^C^6+^ ions, 30keVμm^-1^) at a dose of 150 Gy in the RI-beam factory (RIKEN, Saitama, Japan). The mutant strain 13C3-97, which flowers 14 days earlier, was isolated in M_2_ plants grown in a rice paddy field at the Tohoku University Agricultural Experiment Station in Kashimadai, Osaki City, Miyagi Prefecture (38.46°N, 141.10°E) in 2016. Heading date observation was conducted under controlled SD and LD. The SD condition was 10 hr light (28 ℃)/14 hr dark (24 ℃), and the LD condition was 14.5 hr light (28 ℃) /and 9.5 hr dark (24 ℃). The light condition was provided by white, fluorescent light (approximately 427 μmol m^−2^ s^−1^, on average plant height). The date when the first panicle emerged was recorded as the heading date. The Photoperiod Sensitivity index was defined as PSI = [DTH_LD_ –DTH_SD_]/DTH_LD_ (DTH_LD_ and DTH_SD_ represent days to heading under LD and SD conditions, respectively) (Immark et al. 1997). The BC_1_F_2_ (F_2_) population was derived by crossing 13C3-97 with Nipponbare and self-fertilization. The jmj-M mutant line was selected from the BC_1_F_2_ population as a line carrying a homozygous mutation in the *JMJ706*. The primer sequences used for genotyping the JMJ706 gene were as follows: F: 5’-TGCTTCTGTCCCACAAACTGTTA-3’. R: 5’-AAGTTGCATCCACACAACACATG-3’. Sanger sequencing was performed using BigDyeTM Terminator v 3.1 Cycle sequencing kit (Thermo Fisher Scientific, Massachusetts, USA) and ABI 3730xl Genetic Analyzer (Thermo Fisher Scientific). Leaf age is the total number of leaves on the main stem recorded at the one-week interval.

### Plasmid construction and plant transformation

The jmj-5 and jmj-6 knockout lines were generated using the CRISPR-Cas9 system. The 20-nt target sequence (5′- CTGGGTTCTGTGGCCAGCAG-3′) was inserted to the guide RNA cloning vector, pU6gRNA-oligo (Mikami et al. 2015) at the BbsI sites. The resulting vector was digested with AscI and PacI and subsequently cloned into the binary vector pZK_gYSA_FFCas9 (Mikami et al. 2015). *Agrobacterium tumefaciens* strain EHA105 was transformed with the resulted selected plasmid by electroporation. Then were selected on YEP medium supplemented with 12.5 μg mL^−1^ rifampicin and 100 μg mL^−1^ spectinomycin. Finally, the Agrobacterium-mediated transformation protocol for Nipponbare calli was performed in accordance with the method of Toki et al. (2006).

### Whole genome sequence (WGS) analysis

The plant materials for WGS analysis were 13C3-97 (M_3_), jmj-5 and jmj-6 (T_2_). Genomic DNA was extracted from the leaf tissues using a DNeasy plant mini kit (QIAGEN, Venlo, Netherlands), with IDTE 1×TE solution pH 8.0 (Integrated DNA Technologies, Iowa, USA) instead of the AE buffer provided in the kit for the DNA elution step. The DNA library was prepared using an NEXTFLEX Rapid DNA-seq kit 2.0 (BIOO Scientific, Texas, USA). Whole genome sequencing was performed using the Hiseq instrument (Illumina, California, USA) in paired-end, 2 × 150-bp mode. Bioinformatics analysis for 13C3-97 used the mutation analysis pipeline (Ichida et al. 2019). In the mutation analysis pipeline, mutations were detected using GATK v3 (McKenna et al. 2010), BcfTools (Li, 2011), Pindel (Ye et al. 2009), Delly (Rausch et al. 2012), and Manta (Chen et al. 2016). For the jmj-5 and jmj-6 samples, bioinformatic analysis was performed using The Galaxy platform (The Galaxy Community, 2024) embaded DeepVariant (Poplin et al. 2018). The genome sequence of the japonica cultivar Nipponbare (Os-Nipponbare-Reference-IRGSP-1.0) (Kawahara et al. 2013) was used as a reference genome sequence. The resulting candidate mutations were visually confirmed using the Integrative Genomics Viewer software (IGV) (Robinson et al. 2011).

### Transcriptome analysis

Two leaf samples (each sample contained an equal amount of leaves of three plants) of Nipponbare and jmj-5 were collected under both SD and LD conditions (eight samples in total). Plants were three weeks old. The total RNA was extracted using the RNeasy Plant Mini Kit (QIAGEN, Venlo, Netherlands). The cDNA library preparation was conducted by using VAHTS Universal V8 RNA-seq Library Prep kit (Nanjing, PRC). An NovaSeq X Plus instrument was used for sequencing and a 2×150 paired-end configuration was adopted according to manufacturer’s instructions. Quality control was conducted using Cutadapt (V1.9.1, phred cutoff: 20, error rate: 0.1, adapter overlap: 1bp, min. length: 75, proportion of N: 0.1) (Martin, 2011) to obtain high-quality data. Clean reads of each biological sample were aligned to the *Oryza sativa* IRGSP-1.0 by using Hisat2 (v2.2.1) (Kim et al. 2015). The HTSeq (v0.6.1) software package (Anders et al. 2015) was used to count the reads mapped to genomic features, and to estimate gene and isoform expression levels. The resulting count matrix was used for DEG analysis using DESeq2 (Love et al. 2014) Bioconductor package, a model based on the negative binomial distribution. The DE genes were identified by adjusted *p*-value < 0.05 and a minimum two-fold change in expression.

### Haplotype analysis

The RAP-DB embedded TASUKE+ browser (URL: https://agrigenome.dna.affrc.go.jp/tasuke/ricegenomes/) was used to obtain the sequences and subpopulation data. A total of 685 accessions were used for haplotype analysis, 409 rice varieties were selected based on its frequency (haplotype group with ≥ 5 samples), position of the variation (only the samples having variations in transcript region) and subpopulations (IND: Indica, TEJ: Temperate Japonica, TRJ: Tropical Japonica, JP: Japonica, AUS: Aus, and ARO: Aromatic). Haplotype analysis and network creation were conducted by using pegas (v1.3) (Paradis, 2010) and ape (v5.8.1) (Paradis and Schliep, 2019) software packages in R.

### Gene expression analysis

The fully emerged uppermost leaves of three week old plants were used for all the gene expression experiments. Leaves from three plants were mixed as a sample. As biological replicates, three or four samples were used for Real-Time Quantitative Reverse Transcription PCR (RT-qPCR). Total RNA was extracted from rice leaves using TRIZOL reagent (Invitrogen, ThermoFisher Scientific, Waltham, MA, USA) according to the manufacturer’s instructions and treated with DNase I: TURBO DNA-free™ Kit (Invitrogen). The cDNA was synthesized using 4 μg of total RNA using ReverTra Ace RT qPCR master mix (TOYOBO, Tokyo, Japan). The RT-qPCR was performed with the TaqMan fast universal PCR master mix (Applied Biosystems) on a LightCycler 480 II (Roche, Basel, Switzerland) or StepOnePlus (Applied Biosystems, Massachusetts, USA) according to the manufacturer’s instructions. Gene-specific primers and TaqMan probe sequences are listed in Supplementary Table S2. A rice ubiquitin gene (*Os02g0161900*) was used for normalization. Normalized data were logarithmically transformed (log10) in figure creation for a better representation of expression dynamics and data fluctuations of related genes.

### Chromatin immunoprecipitation followed by quantitative PCR (ChIP-qPCR)

Chromatin extraction and immunoprecipitation were performed using the EpiQuik Plant ChIP Kit (EpigenTek, New York, USA) according to the manufacturer’s protocol with slight modifications. A total weight of 125 mg of mixed leaf samples was harvested from three seedlings (three weeks old). The leaf samples were rinsed in 20 ml of deionized water and crosslinked in 1% formaldehyde under vacuum. The chromatin was extracted and fragmented to 600 bp by sonication using Covaris S2 (Covaris LLC, Massachusetts, USA). The ChIP was performed using the Anti-Histone H3K9me2 (mAbcam 1220, Abcam, Cambridge, UK). The precipitated and input DNAs were quantified by StepOnePlus (Applied Biosystems) according to the manufacturer’s instructions with gene and region-specific primers listed in Supplementary Table S2 using THUNDERBIRDTM SYBR qPCR Mix.

### Accession numbers

Sequence data from this article can be found in NCBI Sequenced Read Archive under the accession number PRJNA1354510. Transcriptome data can be found in Gene Expression Omnibus (https://www.ncbi.nlm.nih.gov/geo/) server under the accession number GSE318488.

## Supporting information

supplemental Files for Rice Jumonji706 confers the photoperiod sensitivity in rice by distinct regulation Manuscript

## Acknowledgments

We thank the RIKEN Nishina Center and the Center for Nuclear Study, the University of Tokyo for operating RIBF for performing the ion-beam irradiation. We appreciate the technical help the Support Unit for Bio-Material Analysis, RIKEN CBS Research Resources Division provides regarding Sanger sequencing. The bioinformatics analysis was performed using the HOKUSAI-BigWaterfall supercomputing system (RIKEN, Saitama, Japan) under project numbers Q22208 and Q23443. We wish to convey our sincere gratitude to Dr. Shintaro Iwasaki, Mari Mito (RNA Systems Biochemistry Laboratory, RIKEN), Dr. Shino Suzuki and Dr. Kazuaki Amikura (Geobiology and Astrobiology Laboratory, RIKEN) for their technical support in experiments. This project was supported by the Cross-ministerial Strategic Innovation Promotion Program (SIP) “Technologies for creating next-generation agriculture, forestry and fisheries” (funding agency: Bio-oriented Technology Research Advancement Institution, NARO)

## Author contributions

A.D.N., R.M. and T.A. conceptualized and designed the study. A.D.N., R.M., Y.S, Y.H., K.I., T.S. and K.T. performed the experiments. H.I. and A.D.N. performed the bioinformatic analysis. A.D.N prepared the figures and drafted the manuscript. A.D.N., R.M., H.I. and T.A revised the manuscript. T.A. did project administration and funding acquisition. All the authors read and approved the final version of the manuscript.

---

**Supplementary Figure S1.**
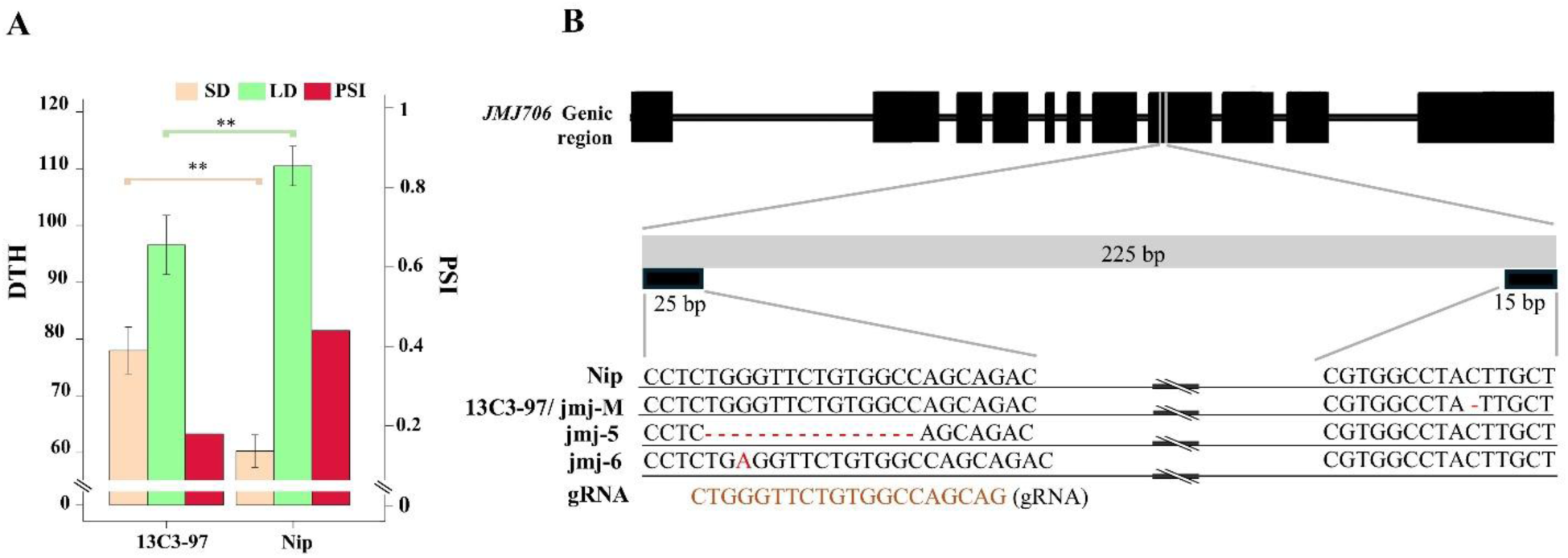
Carbon-ion induced mutant 13C3-97 greatly lost photoperiod sensitivity and nature of the mutations in *JMJ706* mutants. **A**) Days to heading (DTH) and photoperiod sensitivity index (PSI) of mutant 13C3-97 and Nipponbare (Nip) plants under short-day (SD) and long-day (LD) conditions. **B**) Extraction of a nucleotide alignment of Nip, jmj-M, jmj-5 and jmj-6 showing the mutation (red color text) in *JMJ706* gene. Bottom layer sequence is showing the gRNA sequence used for CRISPR-Cas9 constructs. Upper section represents the annotated cDNA sequence and the black color boxes showing the exons.

**Supplementary Figure S2.**
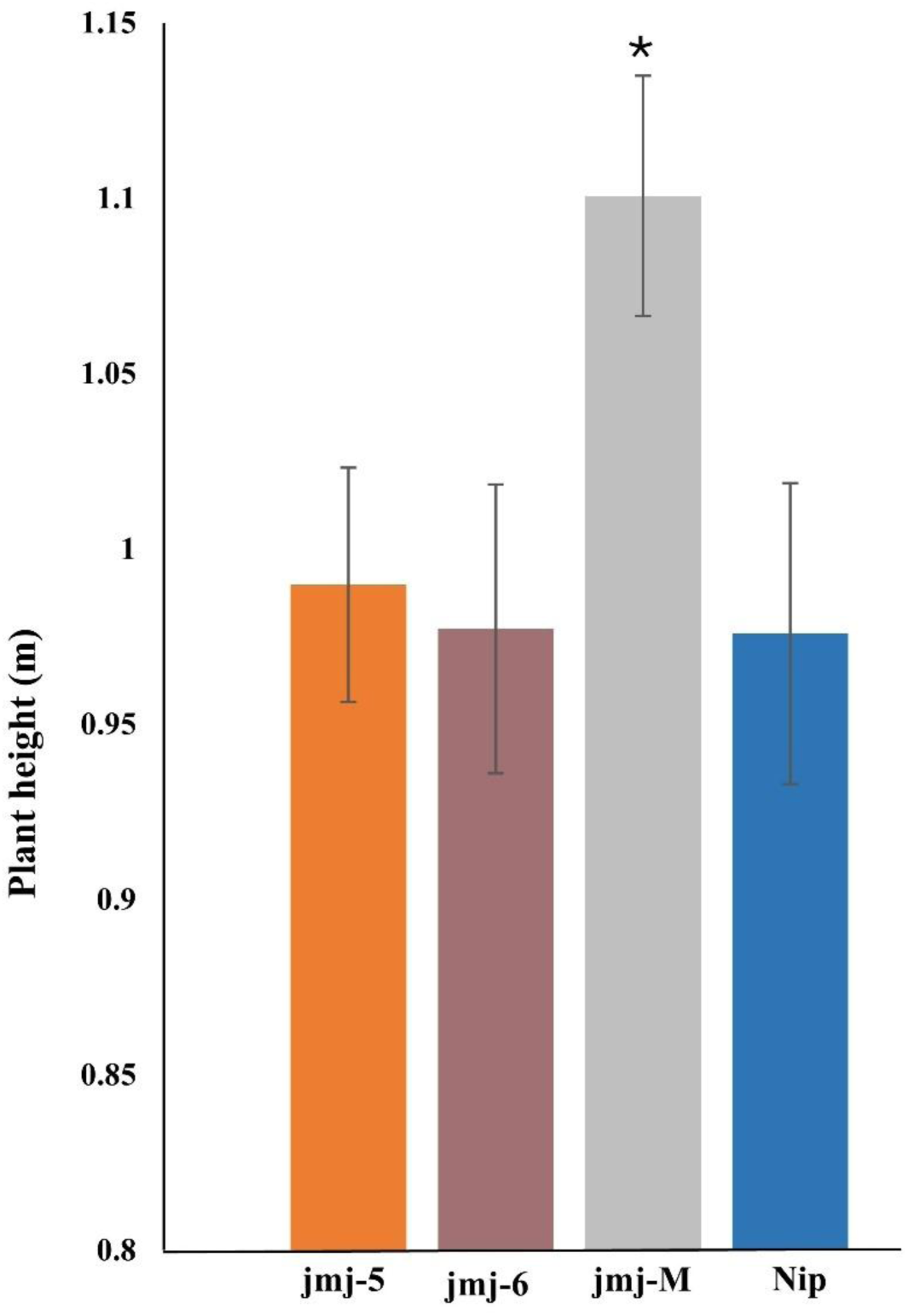
The jmj-M shows higher plant height compared to Nip under SD conditions. Data are means ± S. D. (n>3); and the significance of the difference was assessed by Student’s t-test (*P < 0.05).

**Supplementary Figure S3.**
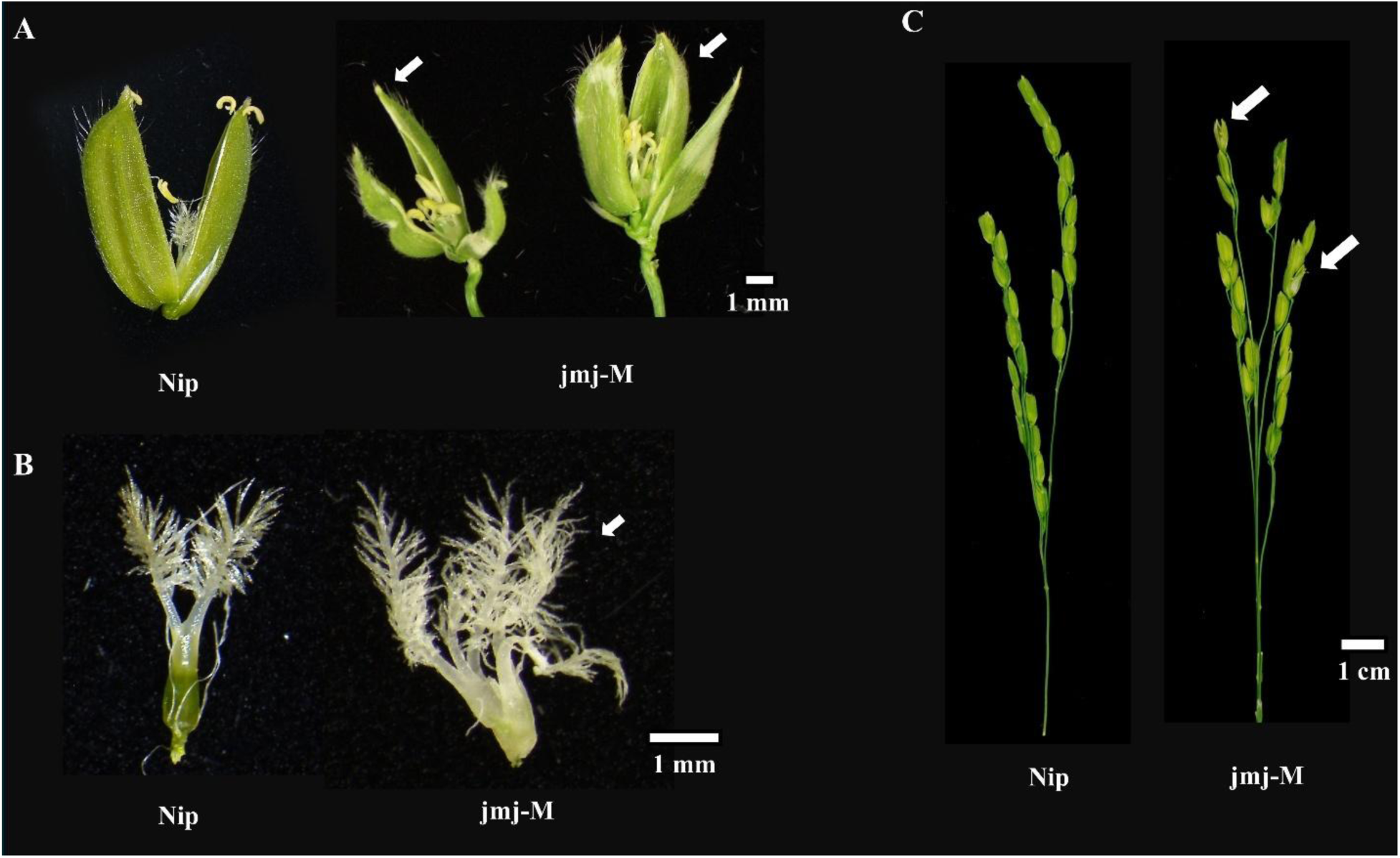
*JMJ706* mutant jmj-M shows defective floral organs compared to Nipponbare (Nip) plants. **A)** Comparison of the lemma and palea between the mutant and Nip. The mutant shows an additional lemma or palea marked with white arrows. **B)** Comparison of the number of pistils between the mutant and Nip. The mutant exhibits several additional pistils (marked with white arrows) compared to Nip. **C)** Comparison of a representative panicle between the mutant and Nip. The mutant panicles show defective floral organs containing spikelets (marked with white arrows). We observed similar defective floral organs in jmj-5 and jmj-6 mutants (data not shown).

## Reference

Ambadas DA, Singh A, Jha RK, Chauhan D, B S, Sharma VK (2023) Genome-wide dissection of AT-hook motif nuclear-localized gene family and their expression profiling for drought and salt stress in rice (*Oryza sativa*). Front Plant Sci 14: 1283555. 10.3389/fpls.2023.1283555

Anders S, Pyl PT, Huber W (2015) HTSeq—a Python framework to work with high-throughput sequencing data. Bioinformatics.15:166–9. 10.1093/bioinformatics/btu638

Andrés F, Galbraith DW, Talón M, Domingo C (2009) Analysis of *PHOTOPERIOD SENSITIVITY5* sheds light on the role of phytochromes in photoperiodic flowering in rice. Plant Physiol 151: 681–690. 10.1104/pp.109.139097

Arora R, Agarwal P, Ray S, Singh AK, Singh VP, Tyagi AK, Kapoor S (2007) MADS-box gene family in rice: genome-wide identification, organization and expression profiling during reproductive development and stress. BMC Genomics 8: 242. 10.1186/1471-2164-8-242

Brambilla V, Fornara F (2013) Molecular control of flowering in response to day length in rice. J Integr Plant Biol 55: 410–418. 10.1111/jipb.12033

Chen X, Hu Y, Zhou D-X (2011) Epigenetic gene regulation by plant Jumonji group of histone demethylase. Biochimica et Biophysica Acta (BBA)-Gene Regulatory Mechanisms 1809: 421–426. 10.1016/j.bbagrm.2011.03.004

Chen X, Schulz-Trieglaff O, Shaw R, Barnes B, Schlesinger F, Källberg M, Cox AJ, Kruglyak S, Saunders CT (2016) Manta: rapid detection of structural variants and indels for germline and cancer sequencing applications. Bioinformatics 32: 1220–1222. 10.1093/bioinformatics/btv710

Chen Z-X, Wu J-G, Ding W-N, Chen H-M, Wu P, Shi C-H (2006) Morphogenesis and molecular basis on naked seed rice, a novel homeotic mutation of *OsMADS1* regulating transcript level of AP3 homologue in rice. Planta 223: 882–890. 10.1007/s00425-005-0141-8

Cheng S, Tan F, Lu Y, Liu X, Li T, Yuan W, Zhao Y, Zhou D-X (2018) WOX11 recruits a histone H3K27me3 demethylase to promote gene expression during shoot development in rice. Nucleic Acids Res 46: 2356–2369. 10.1093/nar/gky017

Corbesier L, Vincent C, Jang S, Fornara F, Fan Q, Searle I, Giakountis A, Farrona S, Gissot L, Turnbull C, et al (2007) FT protein movement contributes to long-distance signaling in floral induction of *Arabidopsis*. Science (1979) 316: 1030–1033. 10.1126/science.1141752

Cui X, Jin P, Cui X, Gu L, Lu Z, Xue Y, Wei L, Qi J, Song X, Luo M (2013) Control of transposon activity by a histone H3K4 demethylase in rice. Proceedings of the National Academy of Sciences 110: 1953–1958. 10.1073/pnas.1217020110

Cui Y, Wang J, Feng L, Liu S, Li J, Qiao W, Song Y, Zhang Z, Cheng Y, Zhang L, et al (2020) A Combination of Long-Day Suppressor Genes Contributes to the Northward Expansion of Rice. Front Plant Sci 11: 864. 10.3389/fpls.2020.00864

Doi K, Izawa T, Fuse T, Yamanouchi U, Kubo T, Shimatani Z, Yano M, Yoshimura A (2004) *Ehd1*, a B-type response regulator in rice, confers short-day promotion of flowering and controls FT-like gene expression independently of *Hd1*. Genes Dev 18: 926–936. 10.1101/gad.1189604

Field A, Adelman K (2020) Evaluating enhancer function and transcription. Annu Rev Biochem 89: 213–234. 10.1146/annurev-biochem-011420-095916

Gao H, Jin M, Zheng X-M, Chen J, Yuan D, Xin Y, Wang M, Huang D, Zhang Z, Zhou K (2014) *Days to heading 7*, a major quantitative locus determining photoperiod sensitivity and regional adaptation in rice. Proceedings of the National Academy of Sciences 111: 16337–16342. 10.1073/pnas.1418204111

Hayama R, Izawa T, Shimamoto K (2002) Isolation of rice genes possibly involved in the photoperiodic control of flowering by a fluorescent differential display method. Plant Cell Physiol 43: 494–504. 10.1093/pcp/pcf059

Hou Y, Wang L, Wang L, Liu L, Li L, Sun L, Rao Q, Zhang J, Huang S (2015) JMJ704 positively regulates rice defense response against Xanthomonas oryzae pv. oryzae infection via reducing H3K4me2/3 associated with negative disease resistance regulators. BMC Plant Biol 15: 286. 10.1186/s12870-015-0674-3

Huang W, Nie H, Feng F, Wang J, Lu K, Fang Z (2019) Altered expression of *OsNPF7. 1* and *OsNPF7. 4* differentially regulates tillering and grain yield in rice. Plant Science 283: 23–31. 10.1016/j.plantsci.2019.01.019

Ichida H, Morita R, Shirakawa Y, Hayashi Y, Abe T (2019) Targeted exome sequencing of unselected heavy-ion beam-irradiated populations reveals less-biased mutation characteristics in the rice genome. The Plant Journal 98: 301–314. 10.1111/tpj.14213

Immark S, Mitchell JM, Jongdee B, Boonwite C, Somrith B, Polvatana A, Fukai S (1997) Determination of phenology development in rainfed lowland rice in Thailand and Lao PDR. Breeding Strategies for Rainfed Lowland Rice in Drought-Prone Environments 77: 89–97.

Ishikawa R, Aoki M, Kurotani KI, Yokoi S, Shinomura T, Takano M, Shimamoto K (2011) Phytochrome B regulates *Heading date 1* (*Hd1*)-mediated expression of rice florigen *Hd3a* and critical day length in rice. Molecular Genetics and Genomics 285: 461–470. 10.1007/s00438-011-0621-4

Itoh H, Nonoue Y, Yano M, Izawa T (2010) A pair of floral regulators sets critical day length for *Hd3a* florigen expression in rice. Nat Genet 42: 635–638. 10.1038/ng.606

Izawa T, Oikawa T, Tokutomi S, Okuno K, Shimamoto K (2000) Phytochromes confer the photoperiodic control of flowering in rice (a short-day plant). Plant Journal 22: 391–399. 10.1046/j.1365-313x.2000.00753.x

Jacob Y, Bergamin E, Donoghue MTA, Mongeon V, LeBlanc C, Voigt P, Underwood CJ, Brunzelle JS, Michaels SD, Reinberg D (2014) Selective methylation of histone H3 variant H3. 1 regulates heterochromatin replication. Science (1979) 343: 1249–1253. 10.1126/science.1248357

Jeon J-S, Lee S, Jung K-H, Yang W-S, Yi G-H, Oh B-G, An G (2000) Production of transgenic rice plants showing reduced heading date and plant height by ectopic expression of rice MADS-box genes. Molecular Breeding 6: 581–592. 10.1023/a:1011388620872

Kawahara Y, de la Bastide M, Hamilton JP, Kanamori H, McCombie WR, Ouyang S, Schwartz DC, Tanaka T, Wu J, Zhou S (2013) Improvement of the *Oryza sativa* Nipponbare reference genome using next generation sequence and optical map data. Rice 6: 1–10. 10.1186/1939-8433-6-4

Kim D, Langmead B, Salzberg SL (2015) HISAT: a fast spliced aligner with low memory requirements. Nat Methods 12: 357–360. 10.1038/nmeth.3317

Kojima S, Takahashi Y, Kobayashi Y, Monna L, Sasaki T, Araki T, Yano M (2002) *Hd3a*, a rice ortholog of the Arabidopsis *FT* gene, promotes transition to flowering downstream of *Hd1* under short-day conditions. Plant Cell Physiol 43: 1096–1105. 10.1093/pcp/pcf156

Komiya R, Ikegami A, Tamaki S, Yokoi S, Shimamoto K (2008) *Hd3a* and *RFT1* are essential for flowering in rice. Development 135: 767–774. 10.1242/dev.008631

Komiya R, Yokoi S, Shimamoto K (2009) A gene network for long-day flowering activates *RFT1* encoding a mobile flowering signal in rice. Development 136: 3443–3450. 10.1242/dev.040170

Le H, Simmons CH, Zhong X (2025) Functions and mechanisms of histone modifications in plants. Annu Rev Plant Biol 76: 551–578. 10.1146/annurev-arplant-083123-070919

Lee D-W, Lee S-K, Rahman MM, Kim Y-J, Zhang D, Jeon J-S (2019) The role of rice vacuolar invertase2 in seed size control. Mol Cells 42: 711–720. 10.14348/molcells.2019.0109

Li H (2011) A statistical framework for SNP calling, mutation discovery, association mapping and population genetical parameter estimation from sequencing data. Bioinformatics 27: 2987–2993. 10.1093/bioinformatics/btr509

Li W, Han Y, Tao F, Chong K (2011) Knockdown of SAMS genes encoding S-adenosyl-l-methionine synthetases causes methylation alterations of DNAs and histones and leads to late flowering in rice. J Plant Physiol 168: 1837–1843. 10.1016/j.jplph.2011.05.020

Li Y, Zhao L, Guo C, Tang M, Lian W, Chen S, Pan Y, Xu X, Luo C, Yi Y (2024) *OsNAC103*, an NAC transcription factor negatively regulates plant height in rice. Planta 259: 35. 10.1007/s00425-023-04309-7

Liu C, Lu F, Cui X, Cao X (2010) Histone methylation in higher plants. Annu Rev Plant Biol 61: 395–420. 10.1146/annurev.arplant.043008.091939

Liu R, Zhao D, Li P, Xia D, Feng Q, Wang L, Wang Y, Shi H, Zhou Y, Chen F (2025) Natural variation in *OsMADS1* transcript splicing affects rice grain thickness and quality by influencing monosaccharide loading to the endosperm. Plant Commun 6: 101178. 10.1016/j.xplc.2024.101178

Love M, Anders S, Huber W (2014) Differential analysis of count data–the DESeq2 package. Genome Biol 15: 10–1186. 10.1186/s13059-014-0550-8

Lu F, Li G, Cui X, Liu C, Wang X, Cao X (2008) Comparative analysis of JmjC domain-containing proteins reveals the potential histone demethylases in Arabidopsis and rice. J Integr Plant Biol 50: 886–896. 10.1111/j.1744-7909.2008.00692.x

Lu M, Li W, Jin L, Zhang Q, Zhu P, Huang J, Hu T (2024) OsNF-YA8 promotes starch accumulation and influences seed traits by positively regulating starch biosynthesis in rice. South African Journal of Botany 171: 85–95. 10.1016/j.sajb.2024.06.004

Martin M (2011) Cutadapt removes adapter sequences from high-throughput sequencing reads. EMBnet journal 17: 10–12. 10.14806/ej.17.1.200

McKenna A, Hanna M, Banks E, Sivachenko A, Cibulskis K, Kernytsky A, Garimella K, Altshuler D, Gabriel S, Daly M (2010) The Genome Analysis Toolkit: a MapReduce framework for analyzing next-generation DNA sequencing data. Genome Res 20: 1297–1303. 10.1101/gr.107524.110

Mikami M, Toki S, Endo M (2015) Comparison of CRISPR/Cas9 expression constructs for efficient targeted mutagenesis in rice. Plant Mol Biol 88: 561–572. 10.1007/s11103-015-0342-x

Nagalla AD, Nishide N, Hibara KI, Izawa T (2021) High ambient temperatures inhibit Ghd7-mediated flowering repression in rice. Plant Cell Physiol, 62: 1745–1759. 10.1093/pcp/pcab129

Nemoto Y, Nonoue Y, Yano M, Izawa T (2016) *Hd1*,a *CONSTANS* ortholog in rice, functions as an Ehd1 repressor through interaction with monocot-specific CCT-domain protein Ghd7. Plant J 86: 221–233. 10.1111/tpj.13168

Osugi A, Itoh H, Ikeda-Kawakatsu K, Takano M, Izawa T (2011) Molecular Dissection of the Roles of Phytochrome in Photoperiodic Flowering in Rice. Plant Physiol 157: 1128–1137. 10.1104/pp.111.181792

Paradis E (2010) pegas: an R package for population genetics with an integrated–modular approach. Bioinformatics 26: 419–420. 10.1093/bioinformatics/btp696

Paradis E, Schliep K (2019) ape 5.0: an environment for modern phylogenetics and evolutionary analyses in R. Bioinformatics 35: 526–528. 10.1093/bioinformatics/bty633

Peng L-T, Shi Z-Y, Li L, Shen G-Z, Zhang J-L (2007) Ectopic expression of *OsLFL1* in rice represses *Ehd1* by binding on its promoter. Biochem Biophys Res Commun 360: 251–256. 10.1016/j.bbrc.2007.06.041

Pfluger J, Wagner D (2007) Histone modifications and dynamic regulation of genome accessibility in plants. Curr Opin Plant Biol 10: 645–652. 10.1016/j.pbi.2007.07.013

Poplin R, Chang P-C, Alexander D, Schwartz S, Colthurst T, Ku A, Newburger D, Dijamco J, Nguyen N, Afshar PT (2018) A universal SNP and small-indel variant caller using deep neural networks. Nat Biotechnol 36: 983–987. 10.1038/nbt.4235

Qi J, Li J, Han X, Li R, Wu J, Yu H, Hu L, Xiao Y, Lu J, Lou Y (2016) Jasmonic acid carboxyl methyltransferase regulates development and herbivory-induced defense response in rice. J Integr Plant Biol 58: 564–576. 10.1111/jipb.12436

Rausch T, Zichner T, Schlattl A, Stütz AM, Benes V, Korbel JO (2012) DELLY: structural variant discovery by integrated paired-end and split-read analysis. Bioinformatics 28: i333–i339. 10.1093/bioinformatics/bts378

Robinson JT, Thorvaldsdóttir H, Winckler W, Guttman M, Lander ES, Getz G, Mesirov JP (2011) Integrative genomics viewer. Nat Biotechnol 29: 24–26. 10.1038/nbt.1754

Sun L, Xu H, Song J, Yang X, Wang X, Liu H, Pang M, Hu Y, Yang Q, Ning X (2024) *OsNAC103*, a NAC transcription factor, positively regulates leaf senescence and plant architecture in rice. Rice 17: 15. 10.1186/s12284-024-00690-3

Sun Q, Zhou D-X (2008) Rice JmjC domain-containing gene *JMJ706* encodes H3K9 demethylase required for floral organ development. Proceedings of the National Academy of Sciences 105: 13679–13684. 10.1073/pnas.0805901105

Takano M, Inagaki N, Xie X, Kiyota S, Baba-Kasai A, Tanabata T, Shinomura T (2009) Phytochromes are the sole photoreceptors for perceiving red/far-red light in rice. Proceedings of the National Academy of Sciences 106: 14705–14710. 10.1073/pnas.0907378106

Takano M, Inagaki N, Xie X, Yuzurihara N, Hihara F, Ishizuka T, Yano M, Nishimura M, Miyao A, Hirochika H, et al (2005) Distinct and cooperative functions of phytochromes A, B, and C in the control of deetiolation and flowering in rice. Plant Cell 17: 3311–3325. 10.1105/tpc.105.035899

The Galaxy Community (2024) The Galaxy platform for accessible, reproducible, and collaborative data analyses: 2024 update. Nucleic Acids Res 52: W83–W94. 10.1093/nar/gkae410

Thomas B, Vince-Prue D (1996) Photoperiodism in plants. Elsevier.

Toki S, Hara N, Ono K, Onodera H, Tagiri A, Oka S, Tanaka H (2006) Early infection of scutellum tissue with Agrobacterium allows high-speed transformation of rice. The Plant Journal 47: 969–976. 10.1111/j.1365-313x.2006.02836.x

Tsuji H, Taoka KI, Shimamoto K (2011) Regulation of flowering in rice: Two florigen genes, a complex gene network, and natural variation. Curr Opin Plant Biol 14: 45–52. 10.1016/j.pbi.2010.08.016

Wang X, Yu Z, Li X, Lu J, Tang Y, Li F, Xu H, Chen W, Xu Q (2025a) *JMJ720* encodes an H3K9me2 demethylase that regulates grain size in rice. The Plant Journal 123: e70462. 10.1111/tpj.70462

Wang X, Yu Z, Xu H, Chen H, Lu J, Li X, Li F, Chen W, Xu Q (2025b) An H3K9me2 demethylase encoded by Jumonji C domain-containing *DT2* regulates drought tolerance in rice. Plant Physiol 198: kiaf280. 10.1093/plphys/kiaf280

Wei X, Xu J, Guo H, Jiang L, Chen S, Yu C, Zhou Z, Hu P, Zhai H, Wan J (2010) *DTH8* suppresses flowering in rice, influencing plant height and yield potential simultaneously. Plant Physiol 153: 1747–1758. 10.1104/pp.110.156943

Williams BP, Gehring M (2020) Principles of epigenetic homeostasis shared between flowering plants and mammals. Trends in Genetics 36: 751–763. 10.1016/j.tig.2020.06.019

Wu W, Zheng X-M, Lu G, Zhong Z, Gao H, Chen L, Wu C, Wang H-J, Wang Q, Zhou K (2013) Association of functional nucleotide polymorphisms at *DTH2* with the northward expansion of rice cultivation in Asia. Proceedings of the National Academy of Sciences 110: 2775–2780. 10.3410/f.717996321.793473988

Xuan H, Shi N, Chen J, Jiang Y, Zhang H, Chu C, Li S, Chen X, Yang H (2024) Physical coupling of H3K4me3 demethylases and Polycomb repressive complex 2 to accelerate flowering in rice. Plant Physiol 195: 1802–1806. 10.1093/plphys/kiae172

Xue W, Xing Y, Weng X, Zhao Y, Tang W, Wang L, Zhou H, Yu S, Xu C, Li X, et al (2008) Natural variation in *Ghd7* is an important regulator of heading date and yield potential in rice. Nat Genet 40: 761–767. 10.1038/ng.143

Yano M, Katayose Y, Ashikari M, Yamanouchi U, Monna L, Fuse T, Baba T, Yamamoto K, Umehara Y, Nagamura Y, et al (2000) *Hd1*, a major photoperiod sensitivity quantitative trait locus in rice, is closely related to the Arabidopsis flowering time gene *CONSTANS*. Plant Cell 12: 2473–2484. 10.2307/3871242

Ye K, Schulz MH, Long Q, Apweiler R, Ning Z (2009) Pindel: a pattern growth approach to detect break points of large deletions and medium sized insertions from paired-end short reads. Bioinformatics 25: 2865–2871. 10.1093/bioinformatics/btp394

Yokoo T, Saito H, Yoshitake Y, Xu Q, Asami T, Tsukiyama T, Teraishi M, Okumoto Y, Tanisaka T (2014) *Se14*, encoding a JmjC domain-containing protein, plays key roles in long-day suppression of rice flowering through the demethylation of H3K4me3 of *RFT1*. PLoS One 9: e96064. 10.1371/journal.pone.0096064

Yoshitake Y, Yokoo T, Saito H, Tsukiyama T, Quan X, Zikihara K, Katsura H, Tokutomi S, Aboshi T, Mori N (2015) The effects of phytochrome-mediated light signals on the developmental acquisition of photoperiod sensitivity in rice. Sci Rep 5: 7709. 10.1038/srep07709

Yu H, Zhang L, He X, Zhang T, Wang C, Lu J, He X, Chen K, Gu W, Cheng S, et al (2022) *OsPHS1* is required for both male and female gamete development in rice. Plant Science 325: 111480. 10.1016/j.plantsci.2022.111480

Yu Z, Wang X, Wang Y, Lu J, Chen H, Li X, Xu H, Li F, Chen W, Xu Q (2025) Epigenetic regulation of *ISPL10* enhances regional adaptability of rice varieties. The Plant Journal 121: e70109. 10.1111/tpj.70109

Zhang J, Zhou X, Yan W, Zhang Z, Lu L, Han Z, Zhao H, Liu H, Song P, Hu Y, et al (2015) Combinations of the *Ghd7*, *Ghd8* and *Hd1* genes largely define the ecogeographical adaptation and yield potential of cultivated rice. New Phytologist 208: 1056–1066. 10.1111/nph.13538

Zhao J, Chen H, Ren D, Tang H, Qiu R, Feng J, Long Y, Niu B, et al (2015a) Genetic interactions between diverged alleles of *Early heading date 1* (*Ehd1*) and *Heading date 3a* (*Hd3a*) / *RICE FLOWERING LOCUS T1* (*RFT1*) control differential heading and contribute to regional adaptation in rice (*Oryza sativa*). New Phytologist 208: 936–948. 10.1111/nph.13503

Zhao L, Tan L, Zhu Z, Xiao L, Xie D (2015b) *PAY1* improves plant architecture and enhances grain yield in rice. The Plant Journal 83: 528–536. 10.1111/tpj.12905

Zhu X, Shen W, Huang J, Zhang T, Zhang X, Cui Y, Sang X, Ling Y, Li Y, Wang N, et al (2018) Mutation of the *OsSAC1* gene, which encodes an endoplasmic reticulum protein with an unknown function, causes sugar accumulation in rice leaves. Plant Cell Physiol 59: 487–499. 10.1093/pcp/pcx203

Zong W, Ren D, Huang M, Sun K, Feng J, Zhao J, Xiao D, Xie W, Liu S, Zhang H, et al (2021) Strong photoperiod sensitivity is controlled by cooperation and competition among *Hd1*, *Ghd7* and *DTH8* in rice heading. New Phytologist 229: 1635–1649. 10.1111/nph.16946

