## supplemental Files for Rice Jumonji706 confers the photoperiod sensitivity in rice by distinct regulation Manuscript for "Rice *Jumonji706* confers the photoperiod sensitivity in rice by distinct regulation of short-day and long-day flowering time regulatory pathways"

Table S1: List of Candidate responsible genes for 13C3-97, based on WGS analysis

| Chromosome | location | Type of the mutation | Gene ID | Annotation | Impact of the mutation |
|---|---|---|---|---|---|
| 07 | 16872763 | SNV | *Os07g0471000* | Endoplasmic reticulum stress sensor | missense |
| 07 | 21234543 | SNV | *Os07g0539200* | Conserved hypothetical protein | missense |
| 08 | 27884712 | SNV | *Os08g0557200* | Metallophosphoesterase domain containing protein | missense |
| 10 | 23029363 | DEL (1bp) | *Os10g0577600* | H3K9 demethylase, Floral organ development | frame shift |

SNV: Single nucleotide variant, DEL: Deletion

Table S2. Primers and probe sequences used in this study

Primers and probes used for RT-qPCR experiments

| **Primer** | **Primer sequence （5'-3'）** | **Taqman Probe** | **Probe sequence** |
|---|---|---|---|
| *Ubq*_RT_Fw | GAGCCTCTGTTCGTCAAGTA | *Ubq*_Probe | TTGTGGTGCTGATGTCTACTTGTGTC |
| *Ubq*_RT_Rw | ACTCGATGGTCCATTAAACC | | |
| *Ghd7*_RT_ Fw | GTACGCGTCCAGAAAAGCCT | *Ghd7*_Probe | TGCCGAGATGAGGCCCCGA |
| *Ghd7*_RT_Rw | TTGGCGAAGCGACCTCTC | | |
| *Hd1*_RT_ Fw | AGCAGCATAGTGGTTATGGAGTTG | *Hd1*_Probe | ACACAGATTCCATCAGCAACAGCATATCTT |
| *Hd1*_RT_Rw | CACCGTGCTGTCTGGTACTATAC | | |
| *Ehd1*_RT_ Fw | GAGGATCGAAGAGCTGAGCA | *Ehd1*_Probe | CATTTGGCAGCACATATTCCGAAAGCA |
| *Ehd1*_RT_Rw | AGGATGACCGGGTTTTTCGA | | |
| *Hd3a*_RT_ Fw | TCTACTTCAACTGCCAGCGC | *Hd3a*_Probe | TCCCGATCGATCTGCTGCATGC |
| *Hd3a*_RT_Rw | TTCAATTGTCTGAACCTGCAATGT | | |
| *RFT1*_RT_ Fw | CAGAACTTCAGCACCAGGAACTT | *RFT1*_Probe | AGCTCTACAACCTCGGCTCGCCG |
| *RFT1*_RT_Rw | TCGCGCTGGCAGTTGA | | |
| *OsGI*_ RT_ Fw | TGAACTCCATCATGAGCCACTAG | *OsGI*_ Probe | AGCTGGAAGTTCCTGCATCTGA |
| *OsGI*_ RT_Rw | TATTCTCCACTCAACATCGGGAC | | |
| *JMJ706*_ RT_ Fw | CGTTCCTGTTTATAAAGCTGTGC | *JMJ706*_ Probe | ACTTTCCCTCGTTCCTACCAC |
| *JMJ706*_ RT_Rw | AGTCACTGATAGCAAAGTTGACA | | |

Primers sequences used in Chip RT-qPCR experiment. The position ID is resemble with the position IDs in Figure 4

| **Position** | **Primer** | **Primer sequence （5'-3'）** | **Position** | **Primer** | **Primer sequence （5'-3'）** |
|---|---|---|---|---|---|
| (I) | (I)_Fw | TGCATATATTATTTTAATACATCTATACAAGTTATA | (q) | (q)_Fw | GAAGCAGATGTAGTCGTTAATTTTTGT |
| | (I)_Rw | AACAAGAGCTGAAAAGCAGGT | | (q)_Rw | AAGTAAATCTTCCATGACTGACAAGTAT |
| (II) | (II)_Fw | CAATGTATATGGGTCTAGTGTGATCTT | (r) | (r)_Fw | GGGTAAAATACCATCTATACAAGACACA |
| | (II)_Rw | CGAATCTTGAGATGAATCTGACGG | | (r)_Rw | TGTATGCATCTTCTATTTAATCCAGAAAAA |
| (III) | (III)_Fw | GTAAGTTAGCTATAGCAGGTGAGGT | (s) | (s)_Fw | TGGTGTAGCCAATGCATCATATAG |
| | (III)_Rw | CCCTGTCTTCTTCTTCTTCTTCTTC | | (s)_Rw | GCCTGTTAGCTGAAACACATTAC |
| (IV) | (IV)_Fw | TGCGTATTCACATTTCATCCTACA | | | |
| | (IV)_Rw | CCTAGGGCAAAATATATTGGGGC | | | |
